# Chimeric chromosome landscapes of human somatic cell cultures show dependence on stress and regulation of genomic repeats by CGGBP1

**DOI:** 10.1101/2021.10.30.466567

**Authors:** Subhamoy Datta, Manthan Patel, Sukesh Kashyap, Divyesh Patel, Umashankar Singh

## Abstract

Genomes of somatic cells in culture are prone to spontaneous mutations due to errors in replication and DNA repair. Some of these errors, such as chromosomal fusions, are not rectifiable and subject to selection or elimination in growing cultures. Somatic cell cultures are thus expected to generate background levels of potentially stable chromosomal chimeras. A description of the landscape of such spontaneously generated chromosomal chimeras in cultured cells will help us understand the factors affecting somatic mosaicism. Here we show that short homology-associated non-homologous chromosomal chimeras occur in normal human fibroblasts and HEK293T cells at genomic repeats. The occurrence of chromosomal chimeras is enhanced by heat stress and depletion of a repeat regulatory protein CGGBP1. We also present evidence of homologous chromosomal chimeras between allelic copies in repeat-rich DNA obtained by methylcytosine immunoprecipitation. The formation of homologous chromosomal chimeras at Alu and L1 repeats increases upon depletion of CGGBP1. Our data are derived from *de novo* sequencing from three different cell lines under different experimental conditions and our chromosomal chimera detection pipeline is applicable to long read as well as short read sequencing platforms. These findings present significant information about the generation, sensitivity and regulation of somatic mosaicism in human cell cultures.

## INTRODUCTION

In somatic cells the randomly occurring mutations create mosaic patterns of different cell clusters representative of different genotypes, including some deleterious ones [1]. The mosaically occurring genomic variants arise out of errors in DNA metabolism, most likely due to the errors in replication and DNA repair. Mutagens and stressors that affect the fidelity of DNA replication and repair accelerate the emergence of genomic variants and add to the mosaicism. This process generates alternative genotypes stochastically and at low frequency. In a population average of mosaic cellular genotypes, these, often single base changes and microsatellite variations, are diluted by the more consistently occurring inherited genotypes [2–4]. The mosaically occurring sequence variants can be segregated from the inherited genotypes by applying different statistical models [5,6]. The confidence in segregating inherited genotypes from mosaically occurring sequence variants can be enhanced by analysing the parental genotypes alongside the offspring’s genotypes. The larger the population doublings the cells go through, the more diverse and complex the somatic mosaicism becomes. Thus, the somatic mosaicism in cultured cells, especially transformed or cancer-derived cell lines, is very difficult to decipher.

Somatic mosaicism, although often described in terms of sequence variants at single base positions, also consists of other kinds of genomic alteration which undergo selection with time [7].

Chromosomal integrity is pivotal to eukaryotic genomes. In sexually reproducing dioecious taxa, each round of gametogenesis and fertilization generates new recombinants within the constraints of the genome volume and structure as defined by a set of homologous chromosome pairs. Gametic recombination and allelic assortment generate genetic diversity that provides the raw material for natural selection. Similarly, mitotically heritable somatic mutations generate diverse subpopulations of cells that undergo clonal evolution. Interestingly, such somatic mutations affect germline inheritance as well [8].

Chromosomal integrity is necessary for normal cellular functioning and chromosomal alterations not rectifiable by the DNA repair pathways are subject to selection or elimination. With some exceptions of natural changes in ploidy, the set of chromosomes normal for a cell is a constraint stabilized through evolution. Healthy development and differentiation depends on maintenance of the ploidy throughout countless cell divisions which is maintained as inherited in the somatic cells. Changes in the gross chromosomal composition, such as loss or gain of chromosome(s) as well as deletion or duplication pose a much larger challenge to cellular survival than single base changes and microsatellite variations [9,10]. Chromosomal rearrangements, unlike base changes, are not rectifiable by the DNA repair mechanisms and can only be eliminated by cell death or adverse effects on cell growth and division. As such, the somatic changes in chromosomal composition lead to erratic cell survival and proliferation and fuel clonal selection of mutant cells and drives tumorigenesis. Many chromosomal rearrangements are in fact an outcome of error-prone repair of double strand breaks. Knowing the spectrum of somatic mosaicism at the level of chromosomal rearrangements is an important element in understanding the genome dynamics through time. The mosaic changes that happen early during development can even be lineage specific. Thus, the longer the somatic age of a cell and more the population doubling of a line, the more complex the somatic genotype landscape is expected to be [11]. Widespread loss of sex chromosomes in ageing somatic cells, especially chromosome X in females and Y in males have been described [12].

Somatically mosaic chromosomal aberrations have been widely described using microarray-based assays that detect copy number variations. Megabase-range deletion and loss of uniparental alleles are more common in somatic tissues than previously appreciated. Copy number polymorphisms (CNPs) can span 100kbs or longer and occur within protein coding genes [13]. CNPs can also include over 2Mb long copy-neutral loss of heterozygosity (LoH), frequently seen in cancer cells and is affected by age [14–17]. Similar chromosomal mosaicism also exists in healthy human tissues and by extension, in freshly isolated and cultured primary cells [18]. The actual mechanisms underlying such chromosomal mosaicism have not been worked out. It is argued that unless inherited through the germline, such changes can occur through DNA repair pathways that involve breakage and ligation of double strand breaks. Other than xenobiotic mutagens that cause DNA strand breaks, what factors (i) accelerate the endogenous DNA damage leading to the strand breaks and (ii) misdirect the repair, remain poorly understood. The occurrence of LoH detected in kilobases to megabases long regions is also attributed only to these logical possibilities of double strand breaks or single strand breaks during replication followed by error-prone end-joining.

Somatic mosaicism is accelerated by environmental stressors [19]. Stressors can cause chromosomal fragmentation [20] and accelerate the large-scale genomic changes. Genome adaptation is a crucial factor in evolution under stresses of the environment [21]. Genomic rearrangements are needed for cell survival when the genome is under stress [22] and macromolecular damage and repair, most importantly that of DNA, guides the influence of stress on macromolecular reorganization, survival and evolution [23]. Stochastic genome alterations in response to environmental stressors is a major contributor to somatic mosaicism and occurs at a rate that is several orders of magnitude higher than that of the somatic point mutations [8,10,24].

Heat stress induces DNA damage through impairment of repair mechanisms that seem to be evolutionarily conserved and it has been observed in cells of a variety of model systems [25–30]. Heat stress induces chromosomal aberration by compromising replication fork progression, impairment of DNA-protein interactions needed for repair and thus heat stressed S-phase cells exhibit more chromosomal instability than those in G1/G2 [29,31–33]. Heat induced DSBs are repaired by homologous recombination (HR) [34] and this process is inherently prone to error that gets exacerbated by heat stress [35–37].

Epigenetic changes can also add to somatic mosaicism [38]. Repeats are prominent sites of homologous recombination in human cells [39–41]. It has been reported that recombination of repeats [42] is mitigated by cytosine methylation [43–46]. In addition to repetitive sequences, the regions with high GC skew and R-loop forming properties are also under tight epigenetic control and have been implicated in genomic instability through promiscuous recombinations [47–50].

Whether there are mechanisms, other than the DNA repair machinery, to mitigate the effects of environmental stressors and epigenetic instability on the somatic mutation load remains unexplored. The high throughput DNA sequencing datasets contain hidden, often ignored, information about the scale and nature of somatic mosaicism. The mosaicism at the level of DNA sequence can be highly variable across different sequencing platforms and hence difficult to establish. However, chromosomal mosaicism can potentially be deciphered with higher confidence. The DNA sequence reads representing the sporadic chromosomal mosaicism events are typically eliminated from data analysis steps as they fail to align to single chromosomes; a necessary condition in most cases. An additional challenge is posed by the presence of repetitive sequences, especially the interspersed repeats, in the DNA sequence reads representing chimeric chromosomes. Our recent works on the human protein CGGBP1 have involved large scale DNA sequence data analyses [51–56]. Since loss of CGGBP1 function has been shown to accelerate chromosomal fusions in cultured fibroblasts [57], the presence of chimeric chromosomal reads in our published sequencing datasets has remained an unexplored possibility. Interestingly, CGGBP1 is also a multifunctional protein [58] with roles in heat stress response [58,59], epigenome homeostasis [52,53,55,58,59], regulation of repetitive sequences [51,53,54] and control of endogenous DNA damage [57].

Here, we have developed a DNA sequence data analysis strategy to reliably detect chimeric chromosomal events with high confidence. Through a step-wise pruning and manual curation of the sequence data we detect chimeric chromosomal events in normal human fibroblasts. We have applied this strategy to find out the effects of heat stress and CGGBP1 depletion on chimeric chromosome occurrence in genomic DNA of normal fibroblasts and HEK293T cells. We show that heat stress and CGGBP1 depletion give rise to chimeric chromosomes at regions with homologous sequences between different chromosomes. These regions are rich in L1 and satellite repeats. By applying a similar strategy based on variant calls in the published as well as newly generated cytosine methylation-enriched DNA sequence datasets, we show that CGGBP1 depletion increases interallelic chimeras as well. Our findings not only shed light on the extent of chromosomal mosaicism prevalent in cell cultures rather also provide some insights into the underlying mechanisms. These findings are important for understanding somatic mosaicism, mutations and loss of heterozygosity.

## MATERIALS AND METHODS

### Cell culture, heat shock, genomic DNA isolation and Nanopore sequencing

Human primary fibroblasts GM02639 (Coriell cell repository) and HEK293T-CT and -KD were cultured in DMEM as described before [51,55].

For the heat stress and recovery experiments using GM02639, the increase in temperature from 37°C to 40°C was done through acclimatization of cells progressively at 38°C, 39°C and 40°C for 24h each. After each round of heat stress, cells were either reverted to 37°C for 24h for recovery before harvesting them for DNA extraction or cultured at a 1°C higher temperature. The temperature for heat stress experiments were established by identifying the maximum tolerated temperature at which these cells could be cultured without any visible loss of cell adhesion and death at 40°C as well as after recovery. Using this method the heat stress temperatures of 41°C and 42°C showed evidence of cell detachment and death upon recovery.

For HEK293T-CT and -KD, the cells were directly subjected to heat stress and at 42°C for 24h the cells could be recovered at 37°C with visible cell death.

Genomic DNA for GM02639, HEK293T-CT and HEK293T-KD were extracted as described before [55]. Briefly, the cells are harvested from (i) two T25 flasks (for GM02639) and (ii) two 100mm dishes (each for HEK293T-CT and -KD at different temperature set points). The cells were lysed using cell lysis buffer (10mM Tris pH 8.0, 100mM NaCl, 25mM EDTA and 0.5% SDS v/v) and 2μl of 10mg/ml Proteinase K (P2308; Sigma). The genomic DNA were isolated using phenol:chloroform:isoamyl alcohol (in 25:24:1 ratio) method followed by ethanol precipitation at −20°C overnight. The DNA was dissolved in nuclease-free water and stored in −20°C.

Nanopore sequencing libraries were prepared using the Ligation Sequencing Kit (SQK-LSK109; Oxford Nanopore Technologies) and sequenced on MinION (Mk1B) using FLO-MIN106 flowcells as per the instructions of the manufacturer. 1μg of DNA was used as input for nick ligation and end repair. Using Agencourt AMPure XP beads, DNA fragments were purified and the manufacturer’s Short Fragment Buffer (SFB) was used to enrich adaptor-ligated DNA fragments of all size ranges. After sequencing adaptor ligation, sequencing was performed for ~48h using real-time base calling using Guppy through MinKnow.

### Methyl(cytosine) DNA immunoprecipitation (MeDIP), sequencing of MeDIP libraries and genomic DNA

MeDIP was performed exactly for GM01391 as described earlier [55]. MeDIP data was already available for GM02639 [55]. The sequencing library for GM01391 and genomic DNA libraries for parents (GM01392 and GM01393) were generated according to the protocol mentioned earlier [51]. The Ion Torrent S5 sequencer was used as the sequencing platform. Using the plug-in “FilterDuplicates” in IonTorrent Suite the sequenced reads were filtered to remove poly-clonals and PCR read-duplicates.

### Sequence data analysis

Sequences were acquired (MinKNOW and Guppy basecaller, ONT), subjected to adapter trimming by Porechop. Sequences from genomic DNA and MeDIP were split into 0.2kb and 0.05kb bins respectively and subjected to end-to-end alignment by *bowtie2*. For sequence and genomic coordinate manipulations, *samtools* and *bedtools* were used. Data was compiled using LibreOffice Spreadsheet and graphs were plotted using deepTools or Prism 9 (GraphPad).

### Interchromosomal chimera detection

Reads of at least 600 bp and 150 bp of length were used to identify chimera in heat stress and MeDIP samples respectively. The reads which had only one 0.2kb and 0.05kb bin unmapped (U bin) were filtered out after the alignment in heat stress or genomic DNA and MeDIP or parental genomic DNA respectively. The flanks of the U bin represent two non-homologous chromosomes (for example chromosomes A and B) such that each flank continues to carry the respective chromosome profile. The A-U-B chimeric reads were further splitted into 0.2kb bins with one base sliding window using *EMBOSS splitter* with the options -size 200 -overlap 199.

### Variant calling and allelic identity establishment

The BAM outputs of the mapped reads were first subjected to generate genotype likelihood individually by *bcftools mpileup*. The variants were called by using *bcftools call* with the options -c and --skip-variants ‘INDELS’. The corresponding records or variants at the same genomic positions were identified in the MeDIP sample in an order (offspring, mother and father) using *bcftools isec* with the options -c all -n +3. The allelic identity for offspring at a given location was established by filtering for the genomic positions and its genotypic information by using *bcftools query*, followed by offsprings’ (GM02639 and GM01391) genotype conferred by only one of the parents.

### TP53BP1 ChIP-sequencing and data analysis

ChIP-sequencing was performed exactly as described earlier [51] using TP53BP1 antibody (NB100-304; Novus Biologicals). Briefly, the cells were cross-linked using 4% formaldehyde solution at 37 °C for 10 min followed by quenching with 125 mM glycine. The cross-linked cells were washed with PBS, harvested and resuspended in an SDS lysis buffer containing 1X protease inhibitor. The cells were lysed on ice for 30 min with intermittent tapping. This was followed by sonication using a Diagenode bioruptor for 30 cycles set at 30s on and 30s off. The mean fragment length was standardised to 150 ± 50 bp. Sonicated lysates were cleared by centrifugation and were incubated overnight at 4°C with antibody-conjugated beads. The beads were washed with IP wash buffers. The cross-links were removed in a reverse cross-linking buffer followed by Proteinase K digestion at 65 °C for 15 min. Reverse cross-linked DNA was purified by DNA purification magnetic beads and used for library preparation and sequencing on the Ion Torrent S5 sequencing platform. The TP53BP1-ChIP sequence reads were filtered for read duplicates and read quality score (Q20) using the built-in options in IonTorrent suite. The quality filtered reads were subjected to end-to-end alignment on hg38. The aligned reads were either subjected to estimate genome-wide distribution at 0.2kb bins using *bedtools coverage* with at least 50% read overlaps or used to calculate the difference of normalised signal from HEK293T-CT to HEK293T-KD using *deepTools bamCompare* with the option *subtract* followed by plotting the signal difference at different repeat flanks genome-wide. The data analysis pipeline for TP53BP1 ChIP-seq is described in the results section.

## RESULTS

### Widespread occurrence of non-homologous chromosomal chimeras in cultured human fibroblasts is enhanced by heat stress

We set out to study the scale and nature of stable somatic mosaicism due to chromosomal fusions in cultured human cells. We sequenced the DNA from a human fibroblast line (Coriell Repository; GM02639, from the skin of a 19 yr old male subject), cultured at prescribed conditions, and established a strategy to detect and describe the non-homologous chromosomal chimeras. Our strategy (Figure 1A) involved the following steps in a series: (i) quality filtering of sequence reads, (ii) elimination of reads that aligned to single chromosomes, (iii) mapping of unaligned reads in non-overlapping fragments of 0.2 kb, (iv) curation of alignments to extract reads composed of fragments uniquely aligning only to two non-homologous chromosomes (for example chromosomes A and B), (v) a single base sliding window unique alignment search to identify the region of transition between the two chromosomes on the chimeric sequence reads (A-U-B where U is the region between the regions aligning to A and B respectively), and (vi) classification of the region U with respect to its similarity to flanking regions of A and B (Figure 1A).

**Figure 1.**
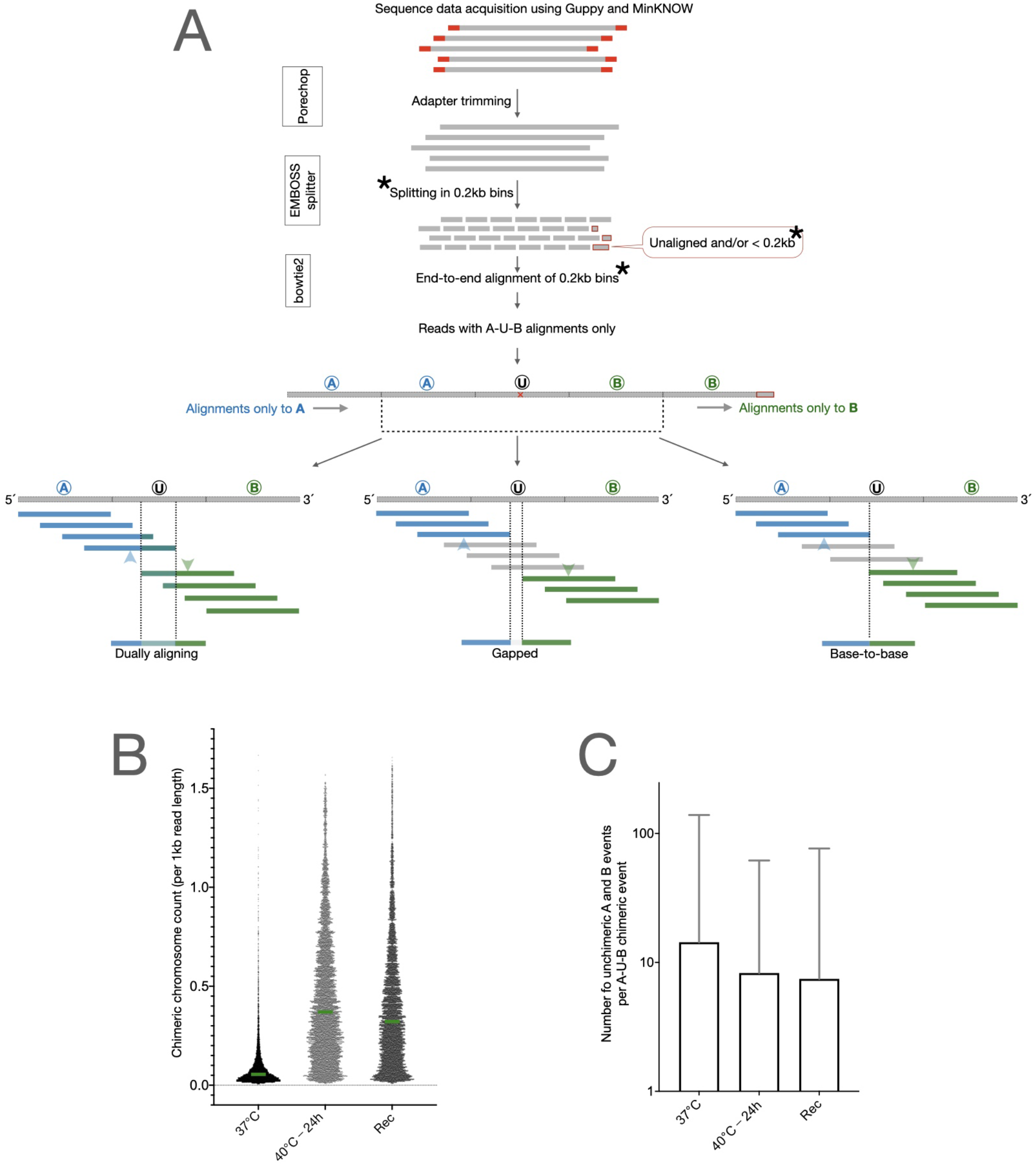
A curated analysis of genomic DNA sequence reads shows evidence of chimeric chromosomal DNA in normal cells that gets enhanced by heat stress: (A) DNA sequence reads were first run through quality check and adapter trimming and then split into end-to-end fragments of 0.2 kb. Analysis was restricted to all the 0.2 kb long read fragments except for the ones at the 3’end of the read which were smaller than 0.2 kb (as all the read-lengths were not multiples of 0.2kb). The 0.2 kb fragments were independently aligned to hg38. All the 0.2 kb fragments of a single reads were expected to align to only one and the same chromosome in hg38. Reads were extracted for which the 0.2 kb fragments aligned to only two different chromosomes with a single 0.2 kb bin serving as the transition point from one chromosome to the other. As shown, these reads were designated as A-U-B reads and the event was termed “occurrence of chimeric chromosomal DNA”. The exact transition point within the 0.2 kb transition fragments were identified by a separate alignment exercise wherein 0.2 kb long fragments were aligned at a time with a single base shift from A towards B such that the shifting fragments alignments to A transitioned to B. Moving from A to B, the last 0.2 kb fragment aligning to A and the first fragment aligning to B are marked by blue and green arrowheads respectively. The various natures of the A-U-B events, dually aligning, base-to-base and gapped, are shown here. (B) The number of chimeric chromosomal DNA events observed in the 37°C sample were enhanced by heat stress in the sample 40°C - 24h and remained increased after recovery in the sample Rec (Unpaired t-test for all three sample pairs, p<0.0001). Green bars show the medians and all data points are plotted. (C) The identification of the chimeric chromosomal DNA events was not an artefact as for a vast majority of A-U-B events there were several non-chimeric A or B alignments observed. The Y axis values are mean±SD.

We discovered that the chimeric chromosomes were prevalent in normal cells at stress-free culture conditions. Only 79.1% of total reads that qualified the quality threshold could be aligned to single chromosomes unambiguously whereas 13.9% of reads represented non-homologous chromosomal chimeras. 7% reads fell out of these two categories and were eliminated from the analysis (Table S1). These cells are expected to have a normal karyotype. We thus interpreted that these non-homologous chromosomal chimeras represented a background load of errors generated sporadically in a small fraction of cells, akin to somatic mosaicism. We could verify that for each non-homologous chromosomal chimeric read detected through this method, there were on average 14.33±124.76 reads (median=2) representing the two chromosomes with no chimeric event (Table 1). The high standard deviation in the number of non-chimeric chromosomal reads for each chimeric event is due to a non-uniform coverage of genomic regions in the sequence data. The background chimeric chromosome frequency (chimeric DNA events per billion bases sequenced) in these fibroblasts at 37°C (see methods), was 1078.69 (Table 1). This translated into just over a single transition to a non-homologous chromosome per 1 million bases sequenced.

**Table 1.** A-U-B non-homologous chimeric events in GM02639.

| Fibroblast (GM02639) sequencing details<br>(data run through Porechop) |  |  |  |
| --- | --- | --- | --- |
| Sample Name | 37°C | 40°C - 24h | Rec |
| Total A-U-B chimeric events | 8633 | 15230 | 15868 |
| Chimeric DNA events per billion bases sequenced | 1078.69 | 3915.97 | 2208.68 |
| Number of non-chimeric reads per chimeric event (mean±SD) | 14.33±124.76 | 8.27±53.45 | 7.45±69.11 |

Non-homologous chromosomal chimeras can occur spontaneously through error-prone non-homologous end-joining, mitotic recombination or a replication template switch between chromosomes. If true, then the occurrence of chromosomal chimeras would be affected by constraining cells to replicate and repair their DNA under conditions of stress. We used a heat shock regimen to study how heat stress affects the chimeric chromosomal landscape. We subjected the fibroblasts to heat shock by sequentially increasing the culture temperature by 1°C every 24 h. Thus, the cells were observed for any visible signs of stress or cell death due to a cumulative exposure to the following temperatures: 38°C, 39°C, 40°C, 41°C and 42°C with a 24 h incubation at each temperature point (see methods for details). We deemed this duration as long enough for the cells to undertake DNA replication and/or repair. At the end of each cumulative heat stress temperature point, cells were also returned to 37°C for 24 h for recovery and resumption of paused DNA replication and repair. When returned to 37°C for a recovery from the varying amounts of heat stress, we first checked if the cells were able to survive or not. The maximum heat stress these cells could tolerate and yet remain adhered during recovery was 40°C for 24 h with only marginal loss of cells (not shown). We sequenced the DNA from two heat stressed samples, (i) 40°C - 24h and (ii) 40°C - 24h followed by recovery at 37°C - 24h, and worked out the non-homologous chimeric chromosome landscape using the strategy described above for the 37°C sample. A comparison of the chimeric chromosomes detected per kilobase of read length in the three samples is shown in figure 1 (Figure 1B). Clearly, the 40°C - 24h heat stress enhanced the occurrence of chromosomal chimeras whereas the 37°C - 24h recovery (Rec) resulted in only a partial reduction. These chimeric chromosome events were sporadic and a majority of chromosomal chimeras were represented by multiple non-chimeric DNA reads (Figure 1C). The non-chimeric reads to chimeric read ratio was decreased upon heat stress and did not reset after recovery (Figure 1C). These results indicated that some processes sensitive to heat stress are involved in the generation of non-homologous chromosomal chimeras. These chromosomal chimeras were also stable and not lost upon recovery from the heat stress.

### CpG-rich L1 and Alu repeats are the sites of spontaneous and heat-induced chromosomal fusions respectively in human fibroblasts

Next we analyzed the sequence properties of the regions of chimeric transitions between different chromosomes (the 0.2kb long region U in the A-U-B chimeric events). The chimeric events were distributed on the different chromosomes as expected from the chromosome sizes (p<0.001 for Pearson r values >0.5 for correlations between all pairs of chromosomal fusions; Figure S1). These regions did not show any significant DNA sequence motif enrichment or overlaps with the chromosomal fragile sites or disease-associated chromosomal fusion sites [60–63] (not shown) To explain if DNA sequence properties predispose certain genomic regions to chromosomal fusions and generate chimeras, we explored the possibility that interspersed repeats could misdirect sequence homology-based DNA repair processes. A RepeatMasker search for repeat contents of the U regions from the chimeras at 37°C showed that they were unusually rich in LINE-1 repeats (40.21% LINE-1 and 6.91% Alu-SINEs). Upon heat shock (40°C - 24h), the chimeras occurred predominantly at LINE-1 but with an increase at Alu SINEs (25.87% LINE-1 and 9.07% Alu-SINEs). The abundance of chimeric events at repeat-free DNA was increased from ~34% at 37°C to ~46% at 40°C - 24h (Table S2). Thus the chromosomes were susceptible to generate chimeras at LINE-1 repeats and heat stress enhanced chimeric events at repeat-free sites and Alu-SINEs.

During the heat stress, the misdirected homology-based strand invasions at Alu-SINE and LINE-1 repeats could facilitate strand annealing between non-homologous chromosomes which would mimic staggered double strand breaks but would not qualify as genuine chimeric DNA. Such strand annealing and staggered double strand breaks could get ligated to generate artefactual chimeric chromosomal DNA during sequencing library preparation that involves DNA end ligation. To establish the stability of the chromosomal chimeras generated upon heat stress, we allowed the cells to recover at 37°C for 24h; a condition sufficient to allow repair of unligated strand invasions. These chimeric chromosomal DNA could not be repaired at 37°C for 24h and were detected post-recovery with the similar fraction of Alu-SINEs and LINE-1 as the heat-stressed sample without any recovery. The recovery simply reduced the chimeric events at repeat-free DNA from 46% to 40% with predominant prevalence of L1 repeats at the chimeric sites (32.41% LINE-1 and 10.55 % Alu-SINEs). Thus, the events we detected as chimeric chromosomal DNA were likely bonafide fusions between two different chromosomal fragments (Table S2).

A subfamily level analysis of the prevalence of Alu-SINEs and LINE-1 showed that the older subfamilies of these repeats were more prone to form chimeras upon heat shock (Table S3). The AluS and L1M subfamilies accounted for the largest increase in chimeric events upon heat stress. The primate-specific LINE-1 subfamily L1P however remained the largest contributor of chimeric events at the U-bins (Table S3).

The abundance of interspersed nuclear elements at the chimeric chromosomal sites posed several possibilities. There could be a crosstalk between the factors that regulate LINE-1 and Alu repeats and the events leading to the formation of chromosomal chimeras. Cytosine methylation and chromatin compaction act as barriers to recombination potential of the interspersed repeats prevalent in the human genome. We have recently described the cytosine methylation landscape in the same fibroblasts as the ones used here for the chimeric DNA identification. CGGBP1 has turned out to be a regulator of cytosine methylation at various subfamilies of Alu and LINE-1 elements [52,53,55]. The 0.2 kb U bins of the chromosomal chimeras showed expected GC-richness (~40%), however, with a more than expected CpG content (>1.25% for all the three samples) (Table S4). We also observed a higher G/C-skew in the U regions of the 40°C - 24h DNA as compared to 37°C DNA (Figure S2). G/C-skew is a property where cytosine methylation is prone to deregulation upon loss of function of CGGBP1 [52] as well as one which facilitates spurious chromosomal recombinations [50].

We next applied the chromosomal chimera detection pipeline to the already published MeDIP datasets from these cells [55]. This provided us with a possibility to answer multiple questions: What is the chimeric DNA landscape in DNA enriched for cytosine methylation and thereby also rich in repeats? Does CGGBP1 depletion affect generation of chromosomal chimera and if so, how? Can the chimeric chromosome detection pipeline be applied to IonTorrent sequencing reads with read lengths ranging around 0.2kb? We first analyzed our recently published MeDIP data and subsequently also generated new datasets in another fibroblast line to verify the findings.

### Chimeric chromosome analysis in MeDIP-DNA from adult human fibroblasts shows a restricted effect of CGGBP1 on chromosome Y-autosome chimeras

The MeDIP-seq data from the fibroblast line (GM02639) was analyzed for evidence of chromosomal fusions. These MeDIP-seq data were generated on the IonTorrent platform with mean read length of approximately 0.2kb using two samples, GM02639-CT and GM02639-KD (GM02639-CT: CGGBP1-non-targeting siRNA; GM02639-KD: CGGBP1-targeting siRNA). We applied the same fusion detection pipeline as described above with one modification: the read fragmentation was done in units of 0.05 kb instead of 0.2 kb (the asterisk-marked step in Figure 1A).

As described for the heat stress experiments, first only the autosomal inter-chromosomal chimeras were analyzed. The chimeric chromosome frequency in GM02639-CT DNA was comparable to those detected in total genomic DNA using ONT platform with no increase observed in GM02639-KD (chimeric DNA events per billion bases sequenced in GM02639-CT and GM02639-KD was 1217.64 and 1727.77 respectively; table 2). The chimeric DNA events in the MeDIP samples showed a different repeat profile compared to the total genomic DNA. The Alu-SINEs repeat content was higher (~20%) than LINE-1 (~12%) and these values remained similar in CT and KD (Table S5). These results showed that acute CGGBP1 depletion by siRNA does not enhance the net rate of chimeric chromosome generation at genomic regions rich in cytosine methylation. We could however apply our chimeric DNA detection strategy to short read sequence datasets (read length range 0.15 to 0.2kb) to detect chromosomal fusions with specificity.

**Table 2.** Details of repeat contents in the U-bins of the non-homologous chimeric events in GM01391-CT and GM01391-KD.

| **Samples** | **GM01391-CT** | **GM01391-KD** |
| --- | --- | --- |
| Reads with chimeric events | 18809 | 30298 |
| Chimeric events per billion bases sequenced | 1217.64 | 1727.77 |
| % Alu-SINEs in reads with chimeric events | 20.21 | 17.44 |
| % LINE-1 in reads with chimeric events | 13.34 | 14.36 |

These published male fibroblast MeDIP datasets were accompanied by sequence data from the parental genomes (GM02640 or paternal and GM02641 or maternal) allowing determination of allelic identities at thousands of locations genome-wide [55]. This provided an opportunity to detect chimeric chromosomal events between two different alleles and its dependence on CGGBP1. To detect the interallelic chimeric events between homologous chromosomes in the MeDIP data, the pipeline was applied with the following additions to the scheme used for non-homologous interchromosomal chimeras: (i) Each uniquely aligned read for the four samples (GM02639-CT, GM02639-KD, Maternal genomic, Paternal genomic) was subject to variant call with respect to the reference hg38. (ii) Variant locations were identified at which the parental genotypes allowed detection of allelic identities of the MeDIP-seq reads. These restricted combinations could be defined as locations where at least one parent had at least one unique allele (identifiable as a DNA sequence). Such unique parent-allele combinations would allow parent of origin annotations of the two alleles at the corresponding genomic locations in the MeDIP DNA from the offspring. (iii) MeDIP-seq reads from the offspring were classified as either maternal, paternal or having a single allelic transition (maternal to paternal or vice versa). (iv) The allelic annotations were curated to eliminate any reads which presented more than one switch between maternal and paternal identities based on the assumption that the read lengths were too small to present two allelic switches. Also, any offspring allelic identities which were different from the parental alleles were considered sporadic mutations or sequencing errors and eliminated. (v) The allelic recombination frequency was calculated as the percentage of total reads with unique parent-allele combinations which showed an allelic switch.

The variant calls, based on which the allelic identities of the reads were established, were mostly single nucleotide variants. Such variants can spontaneously arise in the culture and can interfere with actual parent-of-origin determination of DNA sequence reads. Before calculating the frequency of interallelic chimeric chromosomes, we first needed to establish the somatic mutation rate for CT and KD by segregating the expected genotypes from the unexpected genotypes. The availability of parental (Coriell Repository GM02640 and GM02641) genotypes for GM02639 [55] allowed us to calculate the rate of somatic point mutations.

In the absence of any somatic mutations, the GM02639-CT and -KD genotypes of the offspring are expected to be restricted and predictable by the parental genotypes. All the reads with such genotypes in GM02639-CT and -KD that were not expected from the parental genotypes were regarded as sequence changes due to random somatic mutations in culture, including some sequencing artifacts. Interestingly, over 50% of all the reads at which we could compare GM02639-CT and -KD MeDIP reads with parental reads, we found evidence of unexpected genotypes (Table S6). Assuming that due to the somatic mutations any base (A/T/G/C) occurs on a single allele independently with a 25% probability, the probability of any base mutating to any of the three bases is 75% per allele and about 56% (75% of 75%) on both the alleles at the same location. Thus, the observed ~54% error rate of genotypes at diploid loci (autosomes) was expected. This calculation assumes that MeDIP enriches methylcytosine in the two samples without any net allelic bias. We also made use of the unique case of the maternal-derived X chromosome in the male fibroblasts to verify this mutation frequency. In line with the calculation above, for a monoallelic X chromosome, the observed maternal X genotypes were at nearly 25% of all the X chromosomal genotypes reported in CT and KD MeDIP-seq data (Table S7).

Thus, approximately 50% of the allelic recombinations identified in CT and KD MeDIP DNA would be due to somatic mutations.

The interallelic recombination frequency remained near 4% in both GM02639-CT and GM02639-KD. After correcting for the somatic mutations, the allelic recombination frequencies in these CT and KD were 1.84% and 2% respectively (Table S8). Thus, in these fibroblasts CGGBP1 depletion did not increase interallelic chimeras between homologous chromosomes, just like it had no effect on non-homologous chromosomal chimera. RepeatMasker analysis showed that the Alus or LINEs contents occurred with expected percentage on the reads containing interallelic chimeras and showed no difference between CT and KD (Table S9).

The absence of an allelic counterpart is expected to affect the rate at which the hemizygous chromosomes X and Y generate chimeras with the autosomes. Thus, the chimeras between autosomes and the chromosomes X or Y were analyzed separately. In CT, 5% of total chimeric reads were X-U-autosome (X-U-A) chimeras. In KD this number was observed at 4.77% (Table S10). Unlike chimeras between autosomes and the X chromosome, we observed a clear increase in Y-U-autosome (Y-U-A) chimera frequency upon CGGBP1 knockdown. Whereas 6.9% of reads mapping to the Y chromosome exhibited chimeras with autosomes in CT, in KD the observed value was 9.36% (Table S10).

With these results we concluded that our strategy can be applied to detect inter-allelic chimeric events specifically if parental sequence data is available. CGGBP1 depletion did not affect interchromosomal chimeric events either between non-homologous or homologous chromosomes in DNA enriched for methylcytosine. The GM02639 cells are from a 19 years old male subject. These cells are slow growing and respond to a partial CGGBP1 depletion by exhibiting a growth arrest. To validate these findings further, we needed to perform this analysis in normal cells which grow rapidly and do not exhibit a strong growth arrest upon partial CGGBP1 depletion.

### Chimeric chromosome prevalence in MeDIP-DNA of a female infant’s fibroblast shows a strong dependence on CGGBP1 depletion

The heat stress experiments described above suggested that the chromosomal fusions arise due to cellular growth processes, including replication and DNA repair errors, under stress. The growth arrest and inertness of these cells, especially upon CGGBP1 depletion, could under-represent the changes in chromosomal fusion frequency in KD. Thus, we replicated these experiments in rapidly growing normal fibroblasts that would continue at least some replication and repair of DNA upon CGGBP1 knockdown. We selected a normal fibroblast line from a 9-month old female (Coriell Repository GM01391) for which parental fibroblasts were available as well. These cells showed a much higher rate of proliferation than the GM02639 fibroblasts (not shown). GM01391-CT and GM01391-KD samples were generated in these cells using the same protocols of siRNA transfection followed by MeDIP-seq, as described before (Figure S3). The choice of a female cell line also eliminated the challenges posed by hemizygosity of sex chromosomes in characterization of interchromosomal chimeras.

The non-homologous interchromosomal chimeras in CT were at 1217.64 per billion bases sequenced, which rose 1.4 fold to 1727.77 in KD (Table 2 and table S11). The repeat content of the reads exhibiting chimeric events were 20.21% Alu-SINEs and 13.34% LINE-1 in CT and 17.44% Alu-SINEs and 14.36% LINE-1 in KD (Table 2).

To calculate the fusions between the X chromosome and autosomes, we first needed to establish if our MeDIP-seq actually captured X chromosomal DNA equally in CT and KD. In CT and KD, the fraction of total MeDIP-seq reads aligning to the X chromosome were 4.77% and 2.80% respectively (Table 3). The representational bias against the X chromosome in MeDIP-seq would thus affect the detection of X-U-A chimeric events. We hence calculated the expected X-autosome chimeric events by normalizing the X chromosomal read counts in CT and KD. Against an expected 3.14% (due to a lower capture of X chromosome in KD-MeDIP as compared to CT-MeDIP), we observed that 5.36% of reads mapping to the X chromosome represented chimeras with the autosomes. In CT, the X-U-A chimeric events remained at 7.75% (Table 3). Thus, despite a near 50% decrease in the representation of X chromosome in the MeDIP-seq, the frequency of X-U-A chimeras increased upon CGGBP1 depletion. These findings suggested that in the background of unequal cytosine methylation capture of the X chromosome between CT and KD, there was an increase in the rate at which the X chromosome formed chimeras with autosomes upon CGGBP1 depletion. This effect of differential enrichment of X chromosome in MeDIP-seq however would be inconsequential for interallelic X chromosomal fusion events.

**Table 3.** X-U-A chimeric events in GM01391-CT and GM01391-KD.

| **Samples** | **GM01391-CT** | **GM01391-KD** |
| --- | --- | --- |
| Reads mapped | 67725152 | 80980111 |
| Reads mapped on X chromosome | 3230110 | 2263810 |
| % mapped reads on X chromosome | 4.77 | 2.80 |
| Reads with X-U-A chimera | 1458 | 1625 |
| % X-U-A chimera (observed) | 7.75 | 5.36 |
| % X-U-A chimera (expected) | 7.75 | 3.14 |

The somatic mutation rate was calculated for GM01391 MeDIP-seq data by comparing the CT and KD genotypes with those of the parental cells and calculating the expected and unexpected genotype frequencies. The mutation rates calculated for CT and KD conformed to the same rates as calculated for the son earlier and ranged at 59.08% and 52.66% respectively (Table S12). We observed that upon CGGBP1 depletion, the autosomal interallelic chimeric events were increased significantly. In CT, the interallelic chimeras were detected at 1.68% (0.69% after correction for somatic mutations) of all the reads aligning to the autosomes at locations where the parental genotypes were unique to decipher allelic identities. This increased to 5.29% (2.50% after correction for somatic mutations) in KD (Table 4). Unlike the interchromosomal chimeric reads, the interallelic chimeric reads did not show any unexpected repeat content with no change upon CGGBP1 depletion. Similar to the autosomes, on the X chromosome also we observed a strong increase in interallelic chimeras. The CT interallelic chimera frequency of 1.36% (0.56% after correction for somatic mutations) was increased to 6.13% (2.90% after correction for somatic mutations) in KD (Table 4). The coordinates of the interchromosomal chimeric DNA events are listed in the *GSE169435*.

**Table 4.** Interallelic chimeras detected on autosomes and X chromosomes in GM01391-CT and GM01391-KD.

| **Samples** | **GM01391-CT** | **GM01391-KD** | **GM01391-CT** | **GM01391-KD** |
| --- | --- | --- | --- | --- |
| **Chromosomes** | **Autosomes** |  | **X chromosomes** |  |
| Reads with only maternal allelic identity | 2053554 | 940277 | 91118 | 20649 |
| Reads with only paternal allelic identity | 2376579 | 1002715 | 71182 | 11238 |
| Reads with interallelic chimeras | 75879 | 108513 | 2244 | 2083 |
| Total reads | 4506012 | 2051505 | 164544 | 33970 |
| % Reads with interallelic chimeras | 1.68 | 5.29 | 1.36 | 6.13 |
| % Reads with interallelic chimeras (corrected for somatic mutation rate) | 0.69 | 2.50 | 0.56 | 2.90 |

These results suggested that in a juvenile fibroblast line, the depletion of CGGBP1 indeed enhanced formation of chimeric chromosomes. This enhancement of chromosomal fusions due to CGGBP1 depletion was weaker at inter-chromosomal chimeras, which were associated with more than expected Alu repeats. However, the interallelic chimeras, formed between the homologous chromosomes, were not associated with unexpected amounts of Alu repeats, occurred at a higher frequency and were strongly enhanced by CGGBP1 depletion.

### Inherent resistance against formation of interallelic chimeras within gene bodies is compromised by CGGBP1 depletion

The detection of interallelic chimeras in the MeDIP-seq data was limited only to the regions where the parental genotypes allowed parent-of-origin identification in the offspring. The sporadic nature of these chimeric events is revealed by the following statistics: Overall, the number of non-chimeric reads for every interallelic chimeric event in the male fibroblast was 23.4 for CT and 21.9 for KD. In the female fibroblast however, the overall representation of non-chimeric reads for each interallelic chimeric event was 58.4 in CT and 17.9 in KD (Table S13).

Allelic recombination in somatic cells can be deleterious as they can result in LoH of genes. We next analyzed if the interallelic chimeric events were occurring randomly throughout the genome and to what extent did they occur within the gene bodies. The coverage of the MeDIP-seq reads showing interallelic chimeras within gene bodies was determined. A normalized set of reads (to eliminate any sequencing depth biases between the samples) were randomly selected from each sample for this coverage analysis. The coverage counts were thus obtained for the entire set of genes (UCSC known transcripts) under two conditions: the “observed” wherein the locations of the interallelic chimeric events were as reported by MeDIP-seq analysis, and, “expected” wherein the same locations were randomly reshuffled throughout the genome (Table S14). This expected versus observed analysis showed that, if the allelic chimeras were forming at random locations genome-wide, they would have occurred in over 10K and 14K known transcripts in male and female respectively (Table 5 and table S14). The observed coverage counts for CT and KD in the male fibroblast were 5113 and 3980 respectively. In the female data, the coverage counts were increased from 2757 in CT to 5830 in KD (Table 5 and table S14). These data show that the sporadically occurring allelic recombination is non-random in its genomic localization and it occurs at a rate much lower than randomly expected. There seems to be a paucity of interallelic chimeric events within the known genes. In the female fibroblast, which was more responsive to CGGBP1 depletion, we could observe that depletion of CGGBP1 accelerated the otherwise restrained interallelic chimeric events within gene bodies. The coordinates of the interallelic chimeric DNA events are listed in the *GSE169435*.

**Table 5.**
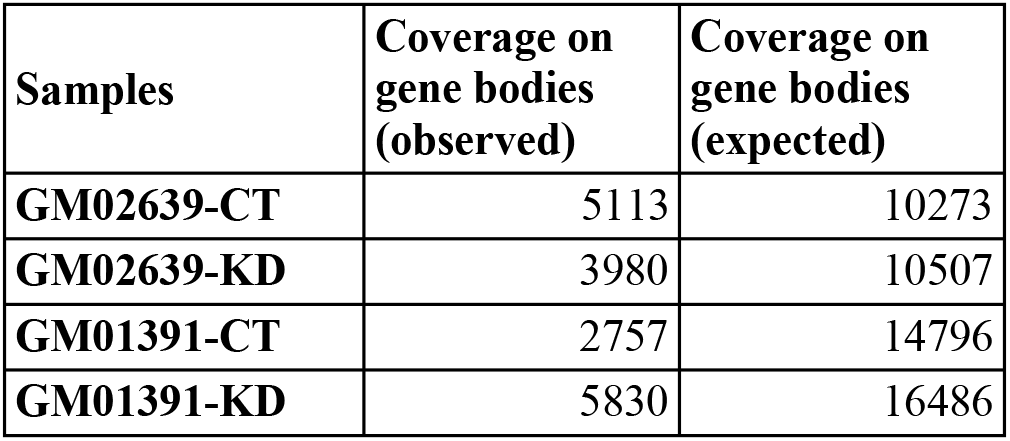
Gene bodies containing interallelic chimeras in GM02639-CT, GM02639-KD, GM01391-CT and GM01391-KD.

These results collectively suggested that chimeric DNA is formed in cultures cells at a background rate that is enhanced by heat stress and CGGBP1 depletion. The MeDIP-seq experiments were conducted in fibroblasts with only a partial (approximately 50%) depletion of CGGBP1. The effect of CGGBP1 depletion was especially pronounced in rapidly growing cells from an infant. The chromatin functions of CGGBP1 have been studied in HEK293T cells. The proliferation of these cells is refractory to CGGBP1 depletion and they make a good system to study the effects of CGGBP1 depletion. We next used the HEK293T stable CT and KD cells for studying the chimeric DNA profiles with or without heat stress.

### Depletion of CGGBP1 mimics the effect of heat stress on chromosomal fusions in HEK293T cells

We have recently reported the effects of CGGBP1 depletion in HEK293T using stable expression of CT (control non-targeting shRNA) and KD (CGGBP1-targeting shRNA). The KD cells have been selected to grow with a >95% knockdown of CGGBP1 [51]. However, the effects of heat stress on chromosomal fusions in these cells is not reported. We first determined the maximum tolerated heat stress for HEK293T-CT and -KD cells. By combining heat stress with HEK293T-CT and -KD we could study how heat stress and CGGBP1 depletion cooperate to generate similar patterns of chromosomal chimeras.

HEK293T-CT and -KD both exhibited high A-U-B type interchromosomal chimeric events per billion bases sequenced (Table 6) at 37°C which were marginally increased in CT and decreased in KD due to heat stress of 42°C for 24h (RM one-way ANOVA on total values of fusions per billion bases yield a P value of 0.0487 and F value 14.87; Figure 2A). Recovery of heat-stressed HEK293T-CT (Rec) and -KD (Rec) samples at 37°C caused significant cell death along with an increase in chimeric events per billion bases sequenced (Figure S4 and table 6). Further analyses of chimeric DNA events in HEK293T-CT and -KD were restricted to 37°C and 42°C - 24h samples only. The chimeric events from these samples showed a genome-wide distribution as expected according to the chromosomal lengths (p<0.001 for Pearson r values >0.5 for correlations between all pairs of chromosomal fusions; Figure S5). There were no obvious regional differences within each chromosome for the chimeric events in CT and KD (Figure S6). The reads representing the chimeric events were rare and embedded within the majority of non-chimeric reads in the same regions (Figure S7).

**Figure 2.**
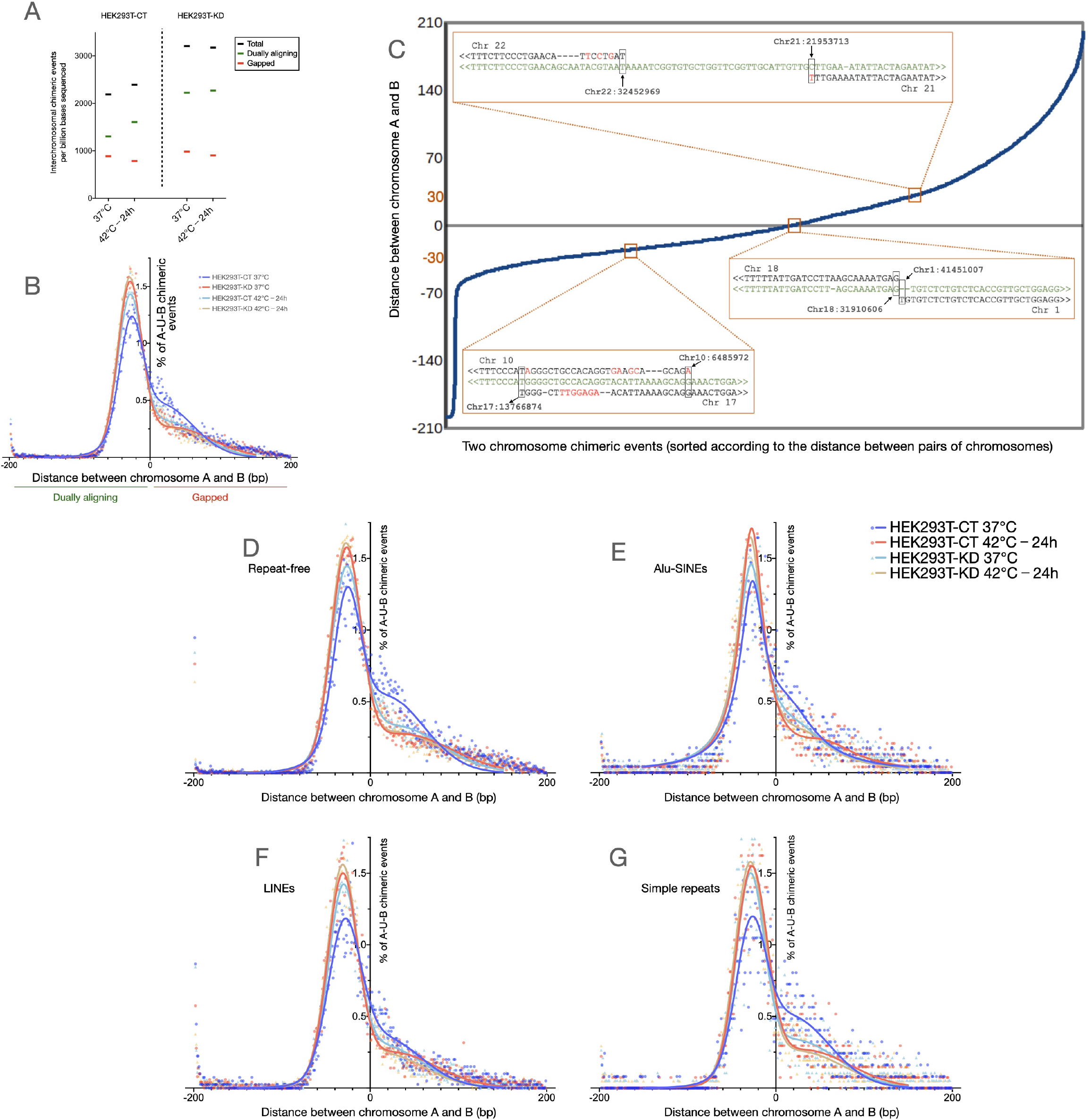
Depletion of CGGBP1 mimics the effects of heat stress on chimeric chromosomal DNA formation in HEK293T cells. (A) The number of chimeric chromosomal DNA events observed per billion bases sequences sequenced in HEK293T cells shows that the chimeric events are enhanced by CGGBP1 depletion as well as heat stress (black data points). The dually aligning chimeric events (green data points) accounted for the increase in the chimeric events in HEK293T-CT 40°C - 24h and HEK293T-KD samples compared to that in the HEK293T-CT 37°C sample. Remarkably, heat stressing the HEK293T-KD sample did not further enhance the chimeric DNA events. The gapped chimeric events (red data points) showed no increase upon heat stress or CGGBP1 depletion. (B) A comparison of the frequency of gapped or dually aligning chimeric events in the same samples as shown in (A). The X-axis shows the gap between the last 0.2 kb fragment of the U bin aligning to A and the first 0.2 kb U bin aligning to B. Negative values depict dually aligning chimeric events, positive values depict gapped events and zero depicts a base-to-base juxtaposition of A and B in the U bin. The Y-axis shows the percent of chimeric events. The first major peak of a double Guassian fit of the frequency distribution shows that the dually aligning events are the least in HEK293T-CT 37°C (blue data points) sample and similarly increased upon heat stress and/or CGGBP1 depletion (aqua, orange and red data points). Such a change was not observed in the gapped alignments. (C) Examples of sequence alignments of the reads at the U bin against the chromosomes A and B. The read sequences are in green, chromosomal genomic sequences are in black with gaps in the alignments shown as “-” and mismatches shown in red. (D-G) The events shown in (B) when split according to repeat contents in their U bin containing the chimeric event show that the effect of CGGBP1 depletion and heat stress in increasing dually aligning chimeric events is maximal at U bins containing no repeats (D) and Alu-SINEs (E) containing U bins show a weak effect of heat stress and/or CGGBP1 depletion on chimeric DNA occurrence. In contrast, the U bins containing LINE (F) and simple repeats (G) showed increased dually aligning chimeric events upon CGGBP1 depletion and/or heat stress.

**Table 6.**
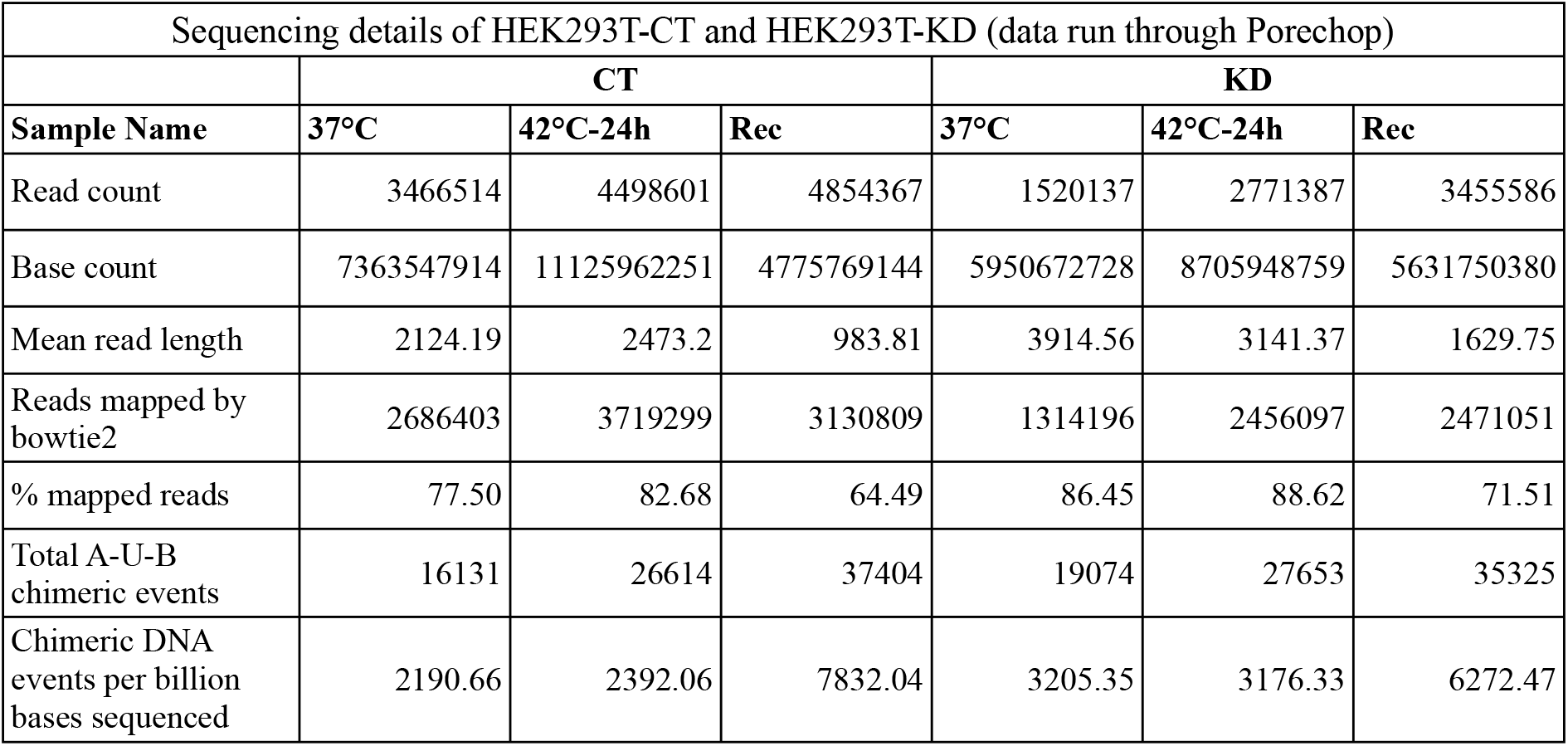
Sequencing details and A-U-B chimeric events at 37°C, 42°C-24h and Rec in HEK293T-CT and HEK293T-KD.

Next, we characterized the nature of these CT and KD chimeric events. The chimeric events were more prevalent in regions with higher G/C-skew (Figure S8). Using a single base window, sliding 5’ to 3’ from chromosome A to B on each A-U-B read, 0.2 kb fragments were generated (Figure 2B) and subjected to alignments. In all the samples the U bins of the majority of chimeric events aligned to both the chromosomes A and B. Most of these dually aligning sequences (Figure 2B) were either exact base-to-base transition points or with short homology (20 to 60 bases long) between A and B. These chimeric events with dually aligning U bins were elevated by heat shock as well as CGGBP1 depletion with only a marginal additive effect of the two treatments (Figure 2B). The less frequent gapped chimeric events, where the U bin sequences did not align to chromosome A or B, did not increase upon heat stress in CT as well as KD (Figure 2B). Representatives of the three different types of chimeric events are shown in figure 2 (Figure 2C). Unlike the short homology events, the frequency of the gapped events were independent of CGGBP1 depletion. For comparison, the short homology events were significantly CGGBP1-dependent (Chi-square 15.78, df 1, z 3.972, P value <0.0001; Figure 2B). The gapped chimeric events showed no significant CGGBP1-dependence (Chi-square 0.4091, df 1, z 0.6396, P value 0.5224).

Short sequence homologies could occur at repeats and so the repetitive sequences could be a target of such short homology chimeric events. We found that heat stress in CT increased short homology chimeric events strongly at LINEs and simple repeats with moderate increases at Alu and repeat-free regions (Figure 2, D to G; ANOVA test details in table S15). Overall, the increases in short homology chimeric events due to CGGBP1 depletion matched (LINEs and simple repeats) or exceeded (Alu and repeat-free sequences) those due to heat stress (Figure 2, D to G). CGGBP1 depletion mimicked the increase in short homology chimeric events caused by heat stress most strongly at LINEs and simple repeats. A combination of CGGBP1 depletion and heat stress did not have an additive effect suggesting that the short homology fusions caused by heat stress involve a deactivation of CGGBP1 (Figure 2, D to G). The coordinates of the fusion events are listed in the GSE169435.

### Enhanced TP53BP1 marks the repeat-rich chromosomal fusion sites induced by CGGBP1 depletion

We have previously described the role of CGGBP1 in heat shock response, endogenous DNA damage and chromosomal fusions independently. Our findings suggested that CGGBP1 depletion and heat stress induce short homology-directed chromosomal rearrangements through overlapping mechanisms that might involve misdirected DNA repair at repeats. TP53BP1, a marker of DNA damage and repair, facilitates recombinational repair [64]. We asked if the formation of short homology chimeric sequences, seemingly caused by CGGBP1 depletion and heat stress through overlapping mechanisms, indeed involved CGGBP1-dependent DNA repair marked by TP53BP1. We performed ChIP-seq for TP53BP1 in CT and KD (sequence data details in table S16). The data analysis pipeline is described in figure S9. The fraction of unaligned reads that could potentially be from repetitive regions and account for chimeric events was higher in KD compared to CT (Table S16). When we extracted the A-U-B events from TP53BP1 ChIP-seq data, we found a staggering amount of Alu-SINEs content (43% in CT and 29% in KD) (Table S17). In CT, TP53BP1 occupancy was restricted to fewer regions with higher coverage per region whereas in KD the TP53BP1 occupancy was spread out with low coverage throughout the genome (Figure 3A). Thus, the focussed and specific TP53BP1 occupancy in CT was disrupted in KD (Figure 3B). Despite a dispersed redistribution of TP53BP1, Alus remained the most prominent repeat content in the A-U-B chimeric events where TP53BP1 was bound in close vicinity (read length 150-200 bps; sonicated input DNA size was 0.3-0.5 kb). These results indicated that in the flanks of Alu of repeats, we might detect an enhanced TP53BP1 occupancy upon CGGBP1 depletion. We measured the difference in TP53BP1 occupancy (normalized CT-KD signals) in flanks of Alu, L1 and simple repeats genome-wide. We observed that in KD, TP53BP1 binding was increased in the flanks of Alu-SINEs, LINEs and simple repeats genome-wide (Figure 3C). These results showed that DNA repair is misdirected to repeat flanks in the absence of CGGBP1 which likely contributes to the short homology fusions that we have observed.

**Figure 3.**
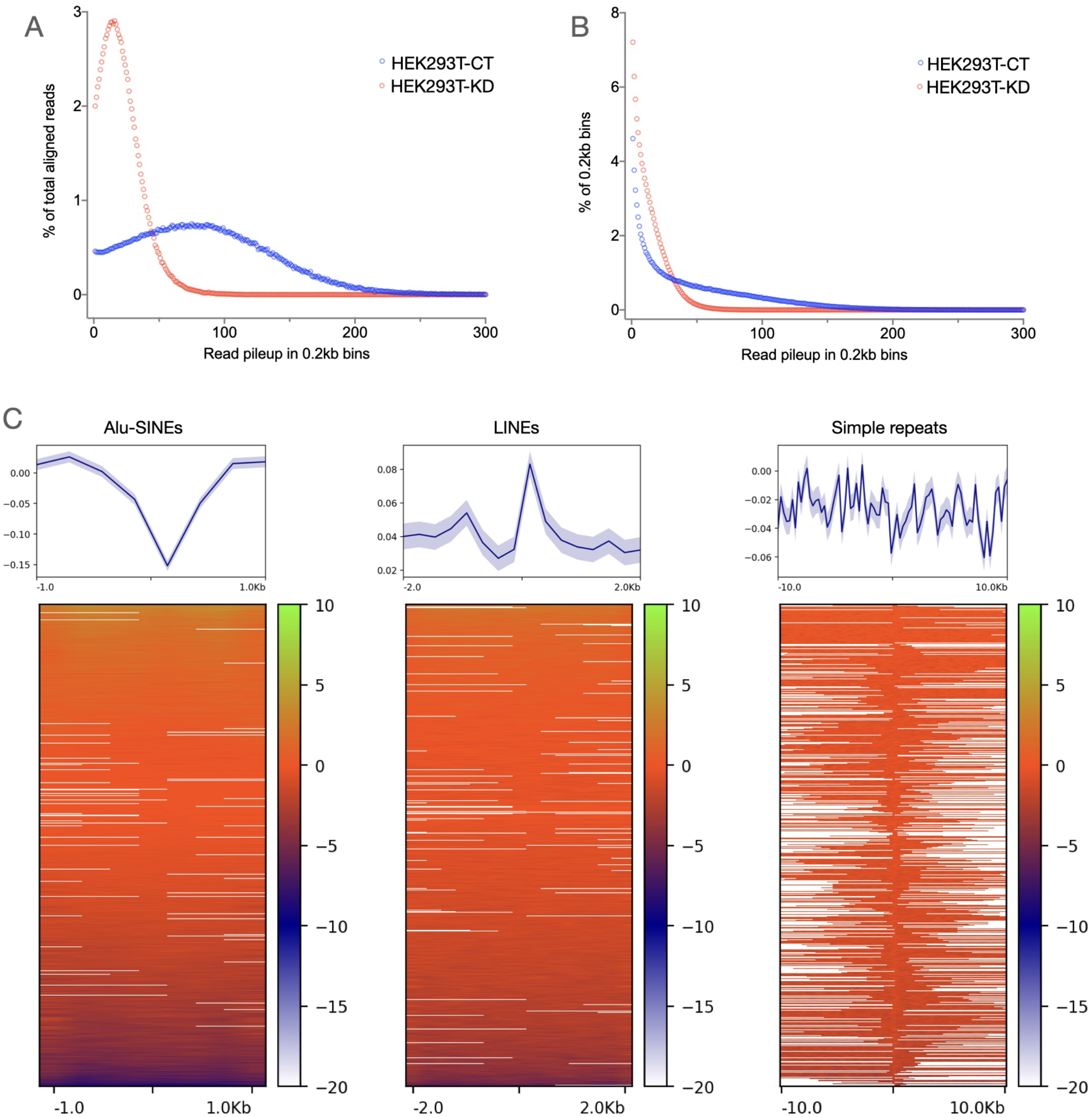
TP53BP1 occupancy change upon CGGBP1 depletion suggests disruption in genomic integrity in the flanks of repeats: (A and B) TP53BP1 ChIP-sequencing reads in HEK293T-CT show a distribution of read pileup peaking around 70-100 (X-axis values for the blue sample in A). Upon CGGBP1 depletion, the peak of read pileup in HEK293T-KD shifts to around 20 (X-axis values for the red sample in A). Accordingly, a majority of HEK293T-KD reads belonged to low read pileups (red sample in B) as compared to HEK293T-CT in which there was a higher prevalence of reads in high read pileup. These results suggest that the TP53BP1 occupancy that is concentrated at a smaller number of genomic sites is diluted and dispersed onto multiple sites genome-wide. (C) The change in TP53BP1 occupancy, calculated as a normalized signal difference HEK293T-CT – HEK293T-KD, shows that TP53BP1 occupancy is enhanced in the flanks of repeat sequences, with the strongest effects observed in the flanks of LINEs and simple repeats, the same repeats at which dually aligning chimeric events are concentrated. The plots in the top panel show mean with standard error.

## DISCUSSION

Somatic mosaicism due to randomly generated mutations is understood to be a fundamental cause for age-related cellular dysfunction and diseases. The net mutation load is a sum of mutations caused by environmental agents and endogenous mechanisms that damage as well as repair the DNA. The alterations in the DNA sequence are rectifiable as the complementary strand serves as a template for BER and NER pathways. The chimeric chromosomal events or chromosomal mosaicism that we have studied here are apparently chromosomal fusion events captured in sequence reads. These chromosomal fusions are formed as an outcome of misdirected double strand repair processes and are not recognized and marked for repair like mismatch or other base changes. These chimeric chromosomes are rare events and can not be determined in the karyotype. The fate of such chimeric chromosomes is dependent on the non-neutrality of their effect on the cells. Thus, these alterations could either persist dormantly or become eliminated with the cells harboring them. Some of these chromosomal chimeras could even undergo clonal selection toward cellular transformation. DNA strand breaks are repaired through homology-directed repair mechanisms or non-homologous end-joining, both of which are error prone.

Alu repeats, the most populous of the repetitive elements in the human genome, present a great challenge to the fidelity of homology mediated repair [65,66]. Alus seed repair directed by sequence microhomology between Alus [67], similar to the short homology chimeric events that we have described. Similarly, L1 repeats induce double strand breaks, possibly through their endonuclease activity, and accelerate homology directed repair or NHEJ [68]. Recombination between Alu and L1 elements have also been reported [69]. Younger Alu subfamilies are reported to generate segmental duplications by homology directed strand break and DNA synthesis [69]. We too observed that in unstressed cells the chromosomal mosaicism was rich mostly in younger Alu and L1 families with the older families showing only an increase after the heat stress.

An estimation of DNA sequence mosaicism is very error-prone. The genuine rare somatic variants could become contaminated with DNA sequence artifacts. However, the chromosomal chimera detection is free from such spurious sequence variations. Our strategy of detection of non-homologous interchromosomal chimeras is based on sequence alignment with length 0.2kb. Single base alterations caused by sequence artifacts would not affect the alignment of such long fragments to target regions in the genome. Additional curation of the data to remove any multiply-aligning reads further ensures that the calls for chimeric chromosomes are free of any errors. There are two attributes of our strategy which strengthen the identification of chimeras: First, the representation of chimeric regions by non-chimeric reads (A-A-A and B-B-B for the A-U-B) ensures that if there were no genuine chimeras, our approach would have classified them as non-chimeric reads. Second, a series of read fragments aligning to chromosomes A and B with a base-by-base incremental identification of the U region is highly unlikely to be artefactual.

The homologous interallelic chimeras can be affected by point mutations. The rigorous curation steps as applied for detecting non-homologous chimeras were applied here as well. However, the MeDIP-seq data were obtained on the IonTorrent platform with shorter read lengths forcing us to use smaller 0.05kb (instead of 0.2kb) fragments for chimera detections. However, even 0.05kb fragments are long enough for high confidence unique alignments and the interchromosomal chimera frequencies calculated using this variation were still comparable to the ones obtained with 0.2kb fragments. IonTorrent base call is of very high confidence [70] and when reinforced with conditions of parental genotypes in the same region and presence of non-chimeric reads representing the chimeric chromosome calls, the errors were minimized further. By comparing the disagreements between the parental and offspring genotypes we were able to work out the mutation (base change) frequency that was used for adjustment of interallelic chimera event rates for the mutation rates. However, the mutation rates were not different between CT and KD and hence would affect both CT and KD equally.

The level of chromosomal mosaicism in cell culture has not been reported before. Fibroblasts and HEK293T are widely used cells. The chromosomal mosaicism in these cells could help us understand the clonal drifts in their populations. This drift can have incalculable consequences on the population-averaged genotypes of cell lines when there are differences in culture conditions that act as stress additives to the background rates of chromosomal chimera formation.

The interallelic chimeras are likely to cause a loss of heterozygosity which would unleash the effects of recessive somatic mutations at heterozygous loci. Since our variant call data are derived from a comparison between parental and offspring DNA sequences, our allelic identification is robust. In the absence of heat stress, the L1 repeats are the primary sites for chromosomal chimera formation with a smaller contribution from the Alu and simple repeats. LINE repeats, a major component of the heterochromatin constitutes a larger fraction of the genome yet remains confined to a smaller volume fraction in the nuclear periphery [71,72]. L1 repeats face a higher molecular crowding [73]. The molecular crowding at the heterochromatin accelerates strand exchange and aberrant repair [74–76]. In growing cells, like DNA replication, DNA repair at the heterochromatic DNA is also delayed [77]. Cytosine methylation at mammalian specific M3 and M4 L1 LINEs is decreased by CGGBP1 depletion [55]. Such a methylation change at L1 repeats can destabilize the genome through aberrant recombinations [43]. Upon heat stress, Alu repeats are transcriptionally activated. The presence of RNA is known to facilitate TP53BP1 association with damage and repair sites. TP53BP1 association with repeats could promote homologous recombination [78].

The results described here are important from multiple perspectives. Through these findings, we get an assessment of the rate at which chromosomal chimera exists in somatic cell cultures. Understanding somatic mutations and mosaicism has advanced our understanding of many diseases including cancer. These findings underscore the importance of the ignored sequence reads in the NGS datasets, often derived from cell cultures under experimental conditions that might be stressful to varying degrees. Changes in somatic mutation profile and scale as an effect of experimental interventions often go unreported. With more and more use of NGS in characterizing the genome and the epigenome, factoring in of such stable spontaneous mutations is key to a complete understanding of the sequence data.

The role of interspersed repeats, most prominently Alu and L1, in endogenous DNA damage is reported but their role in chromosomal mosaicism has not been reported. Our results highlight the role of these repeats in generation of chimeric chromosomes, likely through sequence homology, that then involves DNA damage detection and repair as the repeat flanks are marked by TP53BP1. Because the activity of the repeats are different under conditions of stress, it is reasonable that we detect their differential enrichment in chimeras detected in the heat stressed samples.

Finally, this description of chromosomal mosaicism shows that the protein CGGBP1 levels keep it in check. Depletion of CGGBP1 enhanced the mosaic chromosomal frequency to levels comparable to those of heat stress. CGGBP1 is a gene that shows a heat shock protein-like spike in transcription along with a strong nuclear presence upon acute heat shock. It cooperates with transcription factor NFIX and the high mobility group protein HMGN1 for a proper HSF1 transcription induction upon heat shock. This regulation of heat shock response by CGGBP1 could trick the cells into a heat stress-like state. Thus heat stress and CGGBP1 depletion both could generate similar effects of chromosomal instability thereby inducing strand breaks and repairs though end-ligations. CGGBP1 loss-of-function leads to chromosomal fusions through a mechanism that involves telomere deprotection [57] and the effects mimic breakage-fusion mechanisms of chromosomal fusions typical of cells in crisis. The cells used in this study are low passage primary cells which are not expected to display any chromosomal instabilities. Accordingly, we did not observe any prevalence of telomeric or sub-telomeric repeats in the chimeric reads. The telomeric fusions between the chromosomes are expected to have much longer repeat tracts than the length of our read bins (0.2kb). The fusions observed here however do have a significant contribution of satellite repeats through short sequence homologies across non-homologous chromosomes that might facilitate strand invasions. The other repeats that are enriched in the chimeric DNA identified in this study include the Alu and L1 retrotransposons. As discussed above, these repeats are numerous and strong candidates to generate chimeric chromosomes through short homology-directed end-joining. Interestingly, CGGBP1 regulates both these repeat types. CGGBP1 is required for proper cytosine methylation and inactivation of L1 and Alu repeats. CGGBP1 depletion activates Alu SINEs in a manner that is similar to Alu induction by heat shock. Thus, CGGBP1 depletion and heat shock could increase the Alu-mediated chimera formation through overlapping mechanisms. Similarly, CGGBP1 is required for proper cytosine methylation and H3K9me3 signals at L1 repeats. It is possible that the loss of CTCF-binding at L1 repeats upon CGGBP1 depletion unpacks the chromatin loops thereby easing the strand invasions across different L1 elements. An expected effect of such an epigenetic disruption is generation of chimeric chromosomes in the neighbourhood of L1 repeats. The presence of TP53BP1 in the flanks of these repeats upon CGGBP1 depletion reinforces the idea that a genuine DNA strand break and an impending repair in repeat flanks generates the chimeric DNA that we discover as chromosomal mosaicism. It is likely that additional mechanisms cooperate with stress response and proteins like CGGBP1 to modulate the rate at which chimeric chromosomes emerge in somatic cells.

## DECLARATIONS

### Authors’ contributions

SD, MP, DP performed the experiments, SD, MP, SK analyzed the data, US supervised the work. SD, MP and US wrote the manuscript.

## Acknowledgements

The authors duly acknowledge the Coriell cell repository for the fibroblasts used in this study and NCCS Pune for HEK293T cells.

## Legends

Table S1. GM02639 sequencing details at 37°C, 40°C-24h and Rec.

Table S2. GM02639 repeat-content at 37°C, 40°C-24h and Rec.

Table S3. GM02639 repeat-subfamilies at 37°C, 40°C-24h and Rec.

Table S4. CpG and GC content in U bins of GM02639 at 37°C, 40°C-24h and Rec.

Table S5. GM02639-CT and GM02639-KD repeat-content.

Table S6. MeDIP reads with allelic identities in GM02639-CT and GM02639-KD.

Table S7. X chromosomal allelic identities in GM02639-CT and GM02639-KD.

Table S8. Autosomal allelic identities and interallelic chimeras in GM02639-CT and GM02639-KD.

Table S9. Repeat content in GM02639-CT and GM02639-KD.

Table S10. X-U-A and Y-U-A chimeric events in GM02639-CT and GM02639-KD.

Table S11. Sequencing details of GM01391-CT and GM01391-KD.

Table S12. MeDIP reads with allelic identities in GM01391-CT and GM01391-KD.

Table S13. MeDIP reads with allelic identities and interallelic chimeras in GM02639-CT, GM02639-KD, GM01391-CT and GM01391-KD.

Table S14. Normalized number of MeDIP reads with interallelic chimeras in GM02639-CT, GM02639-KD, GM01391-CT and GM01391-KD. The normalization was done by randomly selecting the same number of reads (before variant calling) from GM02639-CT and GM01391-KD to match the read counts of GM02639-KD, GM01391-CT respectively. The interallelic chimeras were identified again with the normalized read sets.

Table S15. ANOVA test details of repeat types detected in U bins HEK293T-CT and HEK293T-KD at 37°C, 42°C-24h and Rec.

Table S16. TP53BP1 ChIP sequencing details and A-U-B chimeric events in ChIP-sequencing datasets for HEK293T-CT and HEK293T-KD.

Table S17. Repeat content in U bins of TP53BP1 ChIP-seq data from HEK293T-CT and HEK293T-KD.

Figure S1. The dependence of chimeric chromosomal DNA in GM02639 cells on chromosomal lengths indicate the randomness of the events: The expected chimeric events depict the distribution profiles if all the events were distributed evenly on the different autosomes proportional to their lengths. The observed profiles show a strong dependence of chimeric event occurrence on chromosomal lengths. Some exceptions were observed, which were however inconsistent, again indicating no consistent preference for specific interchromosomal chimeric events. For the calculation of these profiles, the A-U-B events were used non-directionally such that the profiles do not represent A-U-B events differently from the B-U-A events.

Figure S2. Regions undergoing chimeric events upon heat stress show high G/C-skew in GM02639: The chimeric events at these higher G/C-skew regions are reparable and hence lost upon recovery post heat stress. The G/C-skew was calculated as (G-C)/(G+C). The data points were fitted to non-linear damped sine wave function with initial decay constant K=4, lambda=0.2 and phase shift=0. The decay constant K values for GM02639 37°C, 40°C - 24h and Rec samples were 10.44, 9.959 and 10.57 respectively.

Figure S3. Depletion of CGGBP1 by siRNA-mediated knockdown in GM01391: A near 50% knockdown of CGGBP1 was achieved by using CGGBP1-targeting siRNA pool. GAPDH is used as a loading control for the amount of protein. Such a partial knockdown of CGGBP1 in the fibroblasts allows studying the cells without a string cell cycle arrest and senescence-like phenotype.

Figure S4. Recovery from heat stress causes high occurrences of chimeric events in HEK293T cells. The total chimeric events in the Rec samples are enhanced to nearly two folds (compared to the respective samples in figure 2A) with a proportionate increase in gapped and dually aligning chimeric events in CT as well as KD. The high levels of chimeric events in these cells were associated with high mortality upon recovery from heat stress.

Figure S5. Similar to GM02639 cells, in HEK293T cells also we observed a dependence of chimeric chromosomal DNA occurrence on chromosomal lengths thereby suggesting a random distribution of the chimeric events without any inter-chromosomal preferences: Like shown in figure S1, the expected chimeric events depict the distribution profiles if all the events were distributed evenly on the different autosomes proportional to their lengths. A strong dependence of chimeric event events on the chromosomal lengths is observed barring some inconsistent exceptions. In this analysis, the A-U-B events were considered non-directional and the A-U-B events were not differentiated from the B-U-A events.

Figure S6. No major intrachromosomal regional differences were observed between the various HEK293T samples in the distribution of chimeric events on the autosomes. The intra-chromosomal distributions of the chimeric events were calculated in a bin length of 5kb. The profile plots in the top panels above the heatmaps have arbitrary units on the Y-axis. The X-axis has a scaled representation of all chromosomes to 0.2 Mb.

Figure S7. The distribution of non-chimeric events at and in the immediate flanks of the chimeric events show that the chimeric events are not artefacts. The Y-axes represent the read count at the chimeric 0.2kb bin centre. The Data are plotted for 0.5kb upstream and downstream. The central enrichment of the non-chimeric reads indicates a non-uniform sequencing coverage in the region.

Figure S8. Regions undergoing chimeric events in HEK293T upon heat stress show high G/C-skew: The chimeric events at these higher G/C-skew regions are reparable and hence lost upon recovery post heat stress. The G/C-skew was calculated as (G-C)/(G+C). The data points were fitted to non-linear damped sine wave function with initial decay constant K=4, lambda=0.2 and phase shift=0. The decay constant K is highest for HEK293T-CT 37°C and lowest for HEK293T-KD 37°C.

Figure S9. Data analysis pipeline for the TP53BP1 ChIP-sequencing. The results of this pipeline are shown in figure 3. The raw data files are available vide GSE169435.

## REFERENCES

1. Fernández LC, Torres M, Real FX. Somatic mosaicism: on the road to cancer. Nature Reviews Cancer. 2016. pp. 43–55. doi:10.1038/nrc.2015.1

2. Mkrtchyan H, Gross M, Hinreiner S, Polytiko A, Manvelyan M, Mrasek K, et al. The human genome puzzle - the role of copy number variation in somatic mosaicism. Curr Genomics. 2010;11: 426–431.

3. Dou Y, Gold HD, Luquette LJ, Park PJ. Detecting Somatic Mutations in Normal Cells. Trends in Genetics. 2018. pp. 545–557. doi:10.1016/j.tig.2018.04.003

4. Thorpe J, Osei-Owusu IA, Avigdor BE, Tupler R, Pevsner J. Mosaicism in Human Health and Disease. Annu Rev Genet. 2020;54: 487–510.

5. Dou Y, Kwon M, Rodin RE, Cortés-Ciriano I, Doan R, Luquette LJ, et al. Accurate detection of mosaic variants in sequencing data without matched controls. Nat Biotechnol. 2020;38: 314–319.

6. Kim B, Won D, Jang M, Kim H, Choi JR, Kim TI, et al. Next-generation sequencing with comprehensive bioinformatics analysis facilitates somatic mosaic APC gene mutation detection in patients with familial adenomatous polyposis. BMC Med Genomics. 2019;12: 103.

7. Narisu N, Rothwell R, Vrtačnik P, Rodríguez S, Didion J, Zöllner S, et al. Analysis of somatic mutations identifies signs of selection during in vitro aging of primary dermal fibroblasts. Aging Cell. 2019;18: e13010.

8. Iourov IY, Vorsanova SG, Yurov YB. Chromosomal mosaicism goes global. Mol Cytogenet. 2008;1: 26.

9. Heng HHQ. Missing heritability and stochastic genome alterations. Nature Reviews Genetics. 2010. pp. 812–812. doi:10.1038/nrg2809-c3

10. Heng HHQ, Liu G, Stevens JB, Abdallah BY, Horne SD, Ye KJ, et al. Karyotype Heterogeneity and Unclassified Chromosomal Abnormalities. Cytogenetic and Genome Research. 2013. pp. 144–157. doi:10.1159/000348682

11. Mkrtchyan H, Gross M, Hinreiner S, Polytiko A, Manvelyan M, Mrasek K, et al. Early embryonic chromosome instability results in stable mosaic pattern in human tissues. PLoS One. 2010;5: e9591.

12. Ouseph MM, Hasserjian RP, Cin PD, Lovitch SB, Steensma DP, Nardi V, et al. Genomic alterations in patients with somatic loss of the Y chromosome as the sole cytogenetic finding in bone marrow cells. Haematologica. 2020. pp. 555–564. doi:10.3324/haematol.2019.240689

13. Sebat J, Lakshmi B, Troge J, Alexander J, Young J, Lundin P, et al. Large-scale copy number polymorphism in the human genome. Science. 2004;305: 525–528.

14. Jacobs KB, Yeager M, Zhou W, Wacholder S, Wang Z, Rodriguez-Santiago B, et al. Detectable clonal mosaicism and its relationship to aging and cancer. Nat Genet. 2012;44: 651–658.

15. Gibson J, Morton NE, Collins A. Extended tracts of homozygosity in outbred human populations. Hum Mol Genet. 2006;15: 789–795.

16. Simon-Sanchez J, Scholz S, Fung H-C, Matarin M, Hernandez D, Gibbs JR, et al. Genome-wide SNP assay reveals structural genomic variation, extended homozygosity and cell-line induced alterations in normal individuals. Hum Mol Genet. 2007;16: 1–14.

17. Žilina O, Koltšina M, Raid R, Kurg A, Tõnisson N, Salumets A. Somatic mosaicism for copy-neutral loss of heterozygosity and DNA copy number variations in the human genome. BMC Genomics. 2015;16: 703.

18. De S. Somatic mosaicism in healthy human tissues. Trends in Genetics. 2011. pp. 217–223. doi:10.1016/j.tig.2011.03.002

19. Vorsanova SG, Yurov YB, Iourov IY. Dynamic nature of somatic chromosomal mosaicism, genetic-environmental interactions and therapeutic opportunities in disease and aging. Mol Cytogenet. 2020;13: 16.

20. Stevens JB, Abdallah BY, Liu G, Ye CJ, Horne SD, Wang G, et al. Diverse system stresses: common mechanisms of chromosome fragmentation. Cell Death Dis. 2011;2: e178.

21. Horne SD, Chowdhury SK, Heng HHQ. Stress, genomic adaptation, and the evolutionary trade-off. Front Genet. 2014;5: 92.

22. Liu G, Stevens J, Horne S, Abdallah B, Ye K, Bremer S, et al. Genome chaos: Survival strategy during crisis. Cell Cycle. 2014. pp. 528–537. doi:10.4161/cc.27378

23. Kültz D. MOLECULAR AND EVOLUTIONARY BASIS OF THE CELLULAR STRESS RESPONSE. Annual Review of Physiology. 2005. pp. 225–257. doi: 10.1146/annurev.physiol.67.040403.103635

24. Heng HHQ. Missing heritability and stochastic genome alterations. Nature reviews. Genetics. 2010.p.813.

25. Henry KA, Blank HM, Hoose SA, Polymenis M. The Unfolded Protein Response Is Not Necessary for the G1/S Transition, but It Is Required for Chromosome Maintenance in Saccharomyces cerevisiae. PLoS ONE. 2010. p. e12732. doi:10.1371/journal.pone.0012732

26. Kantidze OL, Velichko AK, Luzhin AV, Razin SV. Heat Stress-Induced DNA Damage. Acta Naturae. 2016;8: 75.

27. Paul C, Melton DW, Saunders PTK. Do heat stress and deficits in DNA repair pathways have a negative impact on male fertility? Mol Hum Reprod. 2008; 14: 1–8.

28. Thermal Injury Causes DNA Damage and Lethality in Unheated Surrounding Cells: Active Thermal Bystander Effect. J Invest Dermatol. 2010;130: 86–92.

29. Velichko AK, Petrova NV, Kantidze OL, Razin SV. Dual effect of heat shock on DNA replication and genome integrity. Mol Biol Cell. 2012;23: 3450.

30. Tan Z, Chan YJA, Chua YJK, Rutledge SD, Pavelka N, Cimini D, et al. Environmental stresses induce karyotypic instability in colorectal cancer cells. Mol Biol Cell. 2019;30: 42–55.

31. Wang Y, Guan J, Wang H, Wang Y, Leeper D, Iliakis G. Regulation of dna replication after heat shock by replication protein a-nucleolin interactions. J Biol Chem. 2001;276: 20579–20588.

32. Roti Roti JL. Heat-induced alterations of nuclear protein associations and their effects on DNA repair and replication. Int J Hyperthermia. 2007;23: 3–15.

33. Li S, Wu X. Common fragile sites: protection and repair. Cell Biosci. 2020;10: 29.

34. Takahashi A, Mori E, Nakagawa Y, Kajihara A, Kirita T, Pittman DL, et al. Homologous recombination preferentially repairs heat-induced DNA double-strand breaks in mammalian cells. Int J Hyperthermia. 2017;33: 336–342.

35. Guirouilh-Barbat J, Lambert S, Bertrand P, Lopez BS. Is homologous recombination really an error-free process? Front Genet. 2014;0. doi:10.3389/fgene.2014.00175

36. Genet SC, Fujii Y, Maeda J, Kaneko M, Genet MD, Miyagawa K, et al. Hyperthermia inhibits homologous recombination repair and sensitizes cells to ionizing radiation in a time- and temperature-dependent manner. J Cell Physiol. 2013;228: 1473–1481.

37. Laszlo A, Fleischer I. Heat-induced perturbations of DNA damage signaling pathways are modulated by molecular chaperones. Cancer Res. 2009;69: 2042–2049.

38. Youssoufian H, Pyeritz RE. Mechanisms and consequences of somatic mosaicism in humans. Nat Rev Genet. 2002;3: 748–758.

39. McVean G. What drives recombination hotspots to repeat DNA in humans? Philos Trans R Soc Lond B Biol Sci. 2010;365: 1213.

40. Lesecque Y, Glémin S, Lartillot N, Mouchiroud D, Duret L. The red queen model of recombination hotspots evolution in the light of archaic and modern human genomes. PLoS Genet. 2014;10: e1004790.

41. Giordano M, Infantino L, Biggiogera M, Montecucco A, Biamonti G. Heat Shock Affects Mitotic Segregation of Human Chromosomes Bound to Stress-Induced Satellite III RNAs. Int J Mol Sci. 2020;21. doi:10.3390/ijms21082812

42. Yang F, Wang PJ. Multiple LINEs of retrotransposon silencing mechanisms in the mammalian germline. Semin Cell Dev Biol. 2016;59: 118–125.

43. Zamudio N, Barau J, Teissandier A, Walter M, Borsos M, Servant N, et al. DNA methylation restrains transposons from adopting a chromatin signature permissive for meiotic recombination. Genes Dev. 2015;29: 1256–1270.

44. Chen RZ, Pettersson U, Beard C, Jackson-Grusby L, Jaenisch R. DNA hypomethylation leads to elevated mutation rates. Nature. 1998;395: 89–93.

45. Bird A. DNA methylation patterns and epigenetic memory. Genes Dev. 2002;16: 6–21.

46. Goll MG, Bestor TH. Eukaryotic cytosine methyltransferases. Annu Rev Biochem. 2005;74: 481–514.

47. Polak P, Arndt PF. Transcription induces strand-specific mutations at the 5’ end of human genes. Genome Res. 2008;18: 1216–1223.

48. Polak P, Arndt PF. Long-range bidirectional strand asymmetries originate at CpG islands in the human genome. Genome Biol Evol. 2009;1: 189–197.

49. Auton A, Fledel-Alon A, Pfeifer S, Venn O, Ségurel L, Street T, et al. A fine-scale chimpanzee genetic map from population sequencing. Science. 2012;336: 193–198.

50. Aguilera A, García-Muse T. R loops: from transcription byproducts to threats to genome stability. Mol Cell. 2012;46: 115–124.

51. Patel D, Patel M, Datta S, Singh U. CGGBP1 regulates CTCF occupancy at repeats. Epigenetics Chromatin. 2019;12: 57.

52. Patel D, Patel M, Westermark B, Singh U. Dynamic bimodal changes in CpG and non-CpG methylation genome-wide upon CGGBP1 loss-of-function. BMC Res Notes. 2018;11: 419.

53. Agarwal P, Collier P, Fritz MH-Y, Benes V, Wiklund HJ, Westermark B, et al. CGGBP1 mitigates cytosine methylation at repetitive DNA sequences. BMC Genomics. 2015;16: 390.

54. Agarwal P, Enroth S, Teichmann M, Jernberg Wiklund H, Smit A, Westermark B, et al. Growth signals employ CGGBP1 to suppress transcription of Alu-SINEs. Cell Cycle. 2016;15: 1558–1571.

55. Patel M, Patel D, Datta S, Singh U. CGGBP1-regulated cytosine methylation at CTCF-binding motifs resists stochasticity. BMC Genet. 2020;21: 84.

56. Patel D, Patel M, Datta S, Singh U. CGGBP1-dependent CTCF-binding sites restrict ectopic transcription. Cell Cycle. 2021; 1–15.

57. Singh U, Maturi V, Jones RE, Paulsson Y, Baird DM, Westermark B. CGGBP1 phosphorylation constitutes a telomere-protection signal. Cell Cycle. 2014;13: 96–105.

58. Singh U, Westermark B. CGGBP1--an indispensable protein with ubiquitous cytoprotective functions. Ups J Med Sci. 2015;120: 219–232.

59. Singh U, Bongcam-Rudloff E, Westermark B. A DNA sequence directed mutual transcription regulation of HSF1 and NFIX involves novel heat sensitive protein interactions. PLoS One. 2009;4: e5050.

60. Tate JG, Bamford S, Jubb HC, Sondka Z, Beare DM, Bindal N, et al. COSMIC: the Catalogue Of Somatic Mutations In Cancer. Nucleic Acids Res. 2019;47: D941–D947.

61. Hahn Y, Bera TK, Gehlhaus K, Kirsch IR, Pastan IH, Lee B. Finding fusion genes resulting from chromosome rearrangement by analyzing the expressed sequence databases. Proc Natl Acad Sci U S A. 2004;101: 13257–13261.

62. Kim P, Zhou X. FusionGDB: fusion gene annotation DataBase. Nucleic Acids Research. 2019. pp. D994–D1004. doi:10.1093/nar/gky1067

63. Mitelman Database Chromosome Aberrations and Gene Fusions in Cancer. [cited 6 Oct 2021]. Available: https://mitelmandatabase.isb-cgc.org

64. Mirman Z, de Lange T. 53BP1: a DSB escort. Genes Dev. 2020;34: 7–23.

65. Morales ME, White TB, Streva VA, DeFreece CB, Hedges DJ, Deininger PL. The Contribution of Alu Elements to Mutagenic DNA Double-Strand Break Repair. PLoS Genet. 2015;11: e1005016.

66. Kitada K, Aikawa S, Aida S. Alu-Alu Fusion Sequences Identified at Junction Sites of Copy Number Amplified Regions in Cancer Cell Lines. CGR. 2013;139: 1–8.

67. Chromosomal Translocation Mechanisms at Intronic Alu Elements in Mammalian Cells. Mol Cell. 2005;17: 885–894.

68. Gasior SL, Wakeman TP, Xu B, Deininger PL. The human LINE-1 retrotransposon creates DNA double-strand breaks. J Mol Biol. 2006;357: 1383–1393.

69. Beck CR, Garcia-Perez JL, Badge RM, Moran JV. LINE-1 elements in structural variation and disease. Annu Rev Genomics Hum Genet. 2011;12: 187–215.

70. Loman NJ, Misra RV, Dallman TJ, Constantinidou C, Gharbia SE, Wain J, et al. Performance comparison of benchtop high-throughput sequencing platforms. Nat Biotechnol. 2012;30: 434–439.

71. Daban JR. Physical constraints in the condensation of eukaryotic chromosomes. Local concentration of DNA versus linear packing ratio in higher order chromatin structures. Biochemistry. 2000;39: 3861–3866.

72. Bohrmann B, Haider M, Kellenberger E. Concentration evaluation of chromatin in unstained resin-embedded sections by means of low-dose ratio-contrast imaging in STEM. Ultramicroscopy. 1993;49: 235–251.

73. Bancaud A, Huet S, Daigle N, Mozziconacci J, Beaudouin J, Ellenberg J. Molecular crowding affects diffusion and binding of nuclear proteins in heterochromatin and reveals the fractal organization of chromatin. EMBO J. 2009;28: 3785–3798.

74. Feng B, Frykholm K, Nordén B, Westerlund F. DNA strand exchange catalyzed by molecular crowding in PEG solutions. Chem Commun. 2010;46: 8231–8233.

75. Richter K, Nessling M, Lichter P. Experimental evidence for the influence of molecular crowding on nuclear architecture. J Cell Sci. 2007;120: 1673–1680.

76. Cravens SL, Schonhoft JD, Rowland MM, Rodriguez AA, Anderson BG, Stivers JT. Molecular crowding enhances facilitated diffusion of two human DNA glycosylases. Nucleic Acids Res. 2015;43: 4087–4097.

77. Natale F, Scholl A, Rapp A, Yu W, Rausch C, Cardoso MC. DNA replication and repair kinetics of Alu, LINE-1 and satellite III genomic repetitive elements. Epigenetics Chromatin. 2018;11: 61.

78. Tang J, Cho NW, Cui G, Manion EM, Shanbhag NM, Botuyan MV, et al. Acetylation limits 53BP1 association with damaged chromatin to promote homologous recombination. Nat Struct Mol Biol. 2013;20: 317–325.

