## Supplementary material for "Chimeric chromosome landscapes of human somatic cell cultures show dependence on stress and regulation of genomic repeats by CGGBP1": Table S1

| Fibroblast (GM02639) sequencing details (data run through Porechop) | | | |
|---|---|---|---|
| **Sample Name** | **37°C** | **40°C - 24h** | **Rec** |
| Read count | 597103 | 1528595 | 2349238 |
| Base count | 8003216090 | 3889197457 | 7184368217 |
| Mean read length | 13403.41 | 2544.30 | 3058.17 |
| Reads mapped by bowtie2 | 572621 | 1285361 | 2010421 |
| Aligned reads (%) | 95.90 | 84.09 | 85.58 |
