## Supplementary material for "Chimeric chromosome landscapes of human somatic cell cultures show dependence on stress and regulation of genomic repeats by CGGBP1": Table S2

| Repeat types | 37°C | | | 40°C - 24h | | | Rec | | |
|---|---|---|---|---|---|---|---|---|---|
| | Number of elements | Length occupied (bp) | Sequence (%) | Number of elements | Length occupied (bp) | Sequence (%) | Number of elements | Length occupied (bp) | Sequence (%) |
| **SINEs:** | 1142 | 125687 | 7.28 | 2163 | 292474 | 9.6 | 2609 | 347859 | 10.96 |
| **ALUs** | 1067 | 119247 | 6.91 | 1992 | 276179 | 9.07 | 2462 | 334884 | 10.55 |
| **MIRs** | 74 | 6380 | 0.37 | 170 | 16141 | 0.53 | 146 | 12895 | 0.41 |
| **LINEs:** | 4093 | 700490 | 40.57 | 5046 | 800672 | 26.29 | 6567 | 1045142 | 32.93 |
| **LINE1** | 4025 | 694278 | 40.21 | 4928 | 788116 | 25.87 | 6404 | 1028639 | 32.41 |
| **LINE2** | 63 | 5770 | 0.33 | 105 | 11391 | 0.37 | 150 | 15266 | 0.48 |
| **L3/CR1** | 4 | 396 | 0.02 | 9 | 827 | 0.03 | 11 | 1067 | 0.03 |
| **LTR elements:** | 710 | 105263 | 6.1 | 1203 | 164224 | 5.39 | 1260 | 168221 | 5.3 |
| **ERVL** | 54 | 6570 | 0.38 | 149 | 17028 | 0.56 | 129 | 14192 | 0.45 |
| **ERVL-MaLRs** | 166 | 18397 | 1.07 | 397 | 47752 | 1.57 | 406 | 45705 | 1.44 |
| **ERV_classI** | 337 | 54013 | 3.13 | 510 | 75581 | 2.48 | 550 | 81453 | 2.57 |
| **ERV_classII** | 153 | 26283 | 1.52 | 139 | 23087 | 0.76 | 164 | 25964 | 0.82 |
| **DNA elements:** | 149 | 16608 | 0.96 | 330 | 35943 | 1.18 | 343 | 33572 | 1.06 |
| **hAT-Charlie** | 62 | 6633 | 0.38 | 126 | 12964 | 0.43 | 161 | 15661 | 0.49 |
| **TcMar-Tigger** | 58 | 6732 | 0.39 | 142 | 16926 | 0.56 | 133 | 13436 | 0.42 |
| **Unclassified:** | 384 | 51035 | 2.96 | 387 | 41448 | 1.36 | 501 | 55290 | 1.74 |
| **Total interspersed repeats:** | | 999083 | 57.86 | | 1334761 | 43.82 | | 1650084 | 51.99 |
| **Small RNA:** | 174 | 11219 | 0.65 | 261 | 20524 | 0.67 | 266 | 14829 | 0.47 |
| **Satellites:** | 604 | 104211 | 6.04 | 1319 | 223112 | 7.32 | 1179 | 198216 | 6.25 |
| **Simple repeats:** | 571 | 23838 | 1.38 | 1087 | 64160 | 2.11 | 908 | 38319 | 1.21 |
| **Low complexity:** | 56 | 2872 | 0.17 | 128 | 7715 | 0.25 | 105 | 5887 | 0.19 |
| **% of bases masked** | | | 66.1 | | | 54.18 | | | 60.1 |
| **% of bases unmasked** | | | 33.9 | | | 45.82 | | | 39.9 |
