## Supplementary material for "Chimeric chromosome landscapes of human somatic cell cultures show dependence on stress and regulation of genomic repeats by CGGBP1": Table S3

| GM02639 samples | %Alu-SINEs | | | %LINE-1 | | |
|---|---|---|---|---|---|---|
| | AluJ | AluS | AluY | L1H | L1M | L1P |
| 37°C | 9.61 | 48.26 | 42.13 | 2.62 | 10.57 | 86.81 |
| 40°C - 24h | 13.12 | 59.44 | 27.44 | 2.97 | 23.74 | 73.29 |
| Rec | 12.08 | 57.97 | 29.96 | 3.54 | 18.98 | 77.49 |
