## Supplementary material for "Chimeric chromosome landscapes of human somatic cell cultures show dependence on stress and regulation of genomic repeats by CGGBP1": Table S4

| **GM02639 samples** | **37°C** | **40°C - 24h** | **Rec** |
| --- | --- | --- | --- |
| Number of U bins | 8633 | 15230 | 15868 |
| % CpG content | 1.31 | 1.36 | 1.27 |
| % GC content | 41.00 | 41.71 | 40.98 |
