## Supplementary material for "Chimeric chromosome landscapes of human somatic cell cultures show dependence on stress and regulation of genomic repeats by CGGBP1": Table S5

| Repeat types | GM02639-CT | | | GM02639-KD | | |
|---|---|---|---|---|---|---|
| | Number of elements | Length occupied (bp) | Sequence (%) | Number of elements | Length occupied (bp) | Sequence (%) |
| **SINEs:** | 14917 | 1931519 | 20.41 | 6066 | 853679 | 19.58 |
| **ALUs** | 14375 | 1884855 | 19.92 | 5812 | 830905 | 19.06 |
| **MIRs** | 542 | 46664 | 0.49 | 254 | 22774 | 0.52 |
| **LINEs:** | 10123 | 1245583 | 13.16 | 3893 | 509551 | 11.69 |
| **LINE1** | 9441 | 1183032 | 12.5 | 3630 | 484201 | 11.1 |
| **LINE2** | 640 | 58961 | 0.62 | 251 | 24041 | 0.55 |
| **L3/CR1** | 33 | 2655 | 0.03 | 7 | 702 | 0.02 |
| **LTR elements:** | 4130 | 449761 | 4.75 | 1354 | 152978 | 3.51 |
| **ERVL** | 685 | 76136 | 0.8 | 212 | 22163 | 0.51 |
| **ERVL-MaLRs** | 1820 | 203026 | 2.15 | 609 | 71300 | 1.64 |
| **ERV_classI** | 1226 | 132615 | 1.4 | 459 | 51294 | 1.18 |
| **ERV_classII** | 382 | 36316 | 0.38 | 62 | 7004 | 0.16 |
| **DNA elements:** | 1316 | 125318 | 1.32 | 540 | 51774 | 1.19 |
| **hAT-Charlie** | 592 | 52865 | 0.56 | 235 | 20568 | 0.47 |
| **TcMar-Tigger** | 536 | 54842 | 0.58 | 216 | 23943 | 0.55 |
| **Unclassified:** | 1657 | 111204 | 1.18 | 674 | 47300 | 1.08 |
| **Total interspersed repeats:** | | 3863385 | 40.83 | | 1615282 | 37.05 |
| **Small RNA:** | 1572 | 88300 | 0.93 | 628 | 35470 | 0.81 |
| **Satellites:** | 11242 | 2087359 | 22.06 | 6102 | 1164495 | 26.71 |
| **Simple repeats:** | 6888 | 567282 | 6 | 4029 | 356982 | 8.19 |
| **Low complexity:** | 202 | 9405 | 0.1 | 73 | 3522 | 0.08 |
