## Supplementary material for "Chimeric chromosome landscapes of human somatic cell cultures show dependence on stress and regulation of genomic repeats by CGGBP1": Table S6

| **Samples** | **GM02639-CT** | **GM02639-KD** |
|---|---|---|
| Read counts with unexpected allelic identities | 2328783 | 1084232 |
| Read counts with expected allelic identities | 4237592 | 1998307 |
| Somatic mutation rate | 54.96 | 54.26 |
