## Supplementary material for "Chimeric chromosome landscapes of human somatic cell cultures show dependence on stress and regulation of genomic repeats by CGGBP1": Table S7

| **Samples** | **GM02639-CT** | **GM02639-KD** |
|---|---|---|
| X chromosomal read count with maternal allelic identity | 16548 | 7947 |
| Total X chromosomal read count with non-maternal allelic identity | 48160 | 21816 |
| Total X chromosmal read count subjected to allele identification | 64708 | 29763 |
| Somatic mutation rate for X chromosome | 25.57 | 26.70 |
