## Supplementary material for "Chimeric chromosome landscapes of human somatic cell cultures show dependence on stress and regulation of genomic repeats by CGGBP1": Table S8

| Samples | GM02639-CT | GM02639-KD |
|---|---|---|
| Autosomal reads with maternal allelic identity | 816032 | 394787 |
| Autosomal reads with paternal allelic identity | 985983 | 466003 |
| Autosomal reads with chimeric alleleic identity | 76874 | 39370 |
| Total autosomal reads subjected to allelic identification | 1878889 | 900160 |
| Interallelic chimera frequency | 4.09 | 4.37 |
| Somatic mutation rate for autosomes | 54.96 | 54.26 |
| Corrected interallelic chimera frequency | 1.84 | 2.00 |
