## Supplementary material for "Chimeric chromosome landscapes of human somatic cell cultures show dependence on stress and regulation of genomic repeats by CGGBP1": Table S9

| Repeat types | GM02639-CT | | | GM02639-KD | | |
|---|---|---|---|---|---|---|
| | Number of elements | Length occupied (bp) | Sequence (%) | Number of elements | Length occupied (bp) | Sequence (%) |
| **SINEs:** | 4228 | 610798 | 16.54 | 2847 | 421569 | 16.77 |
| **ALUs** | 3761 | 557317 | 15.09 | 2517 | 385113 | 15.32 |
| **MIRs** | 465 | 53291 | 1.44 | 327 | 36130 | 1.44 |
| **LINEs:** | 3897 | 725424 | 19.65 | 2543 | 465694 | 18.52 |
| **LINE1** | 3385 | 656389 | 17.78 | 2178 | 420403 | 16.72 |
| **LINE2** | 467 | 62928 | 1.7 | 332 | 41635 | 1.66 |
| **L3/CR1** | 29 | 4326 | 0.12 | 26 | 3085 | 0.12 |
| **LTR elements:** | 1721 | 298335 | 8.08 | 1108 | 193987 | 7.72 |
| **ERVL** | 312 | 53566 | 1.45 | 193 | 34211 | 1.36 |
| **ERVL-MaLRs** | 686 | 112482 | 3.05 | 467 | 75439 | 3 |
| **ERV_classI** | 612 | 111808 | 3.03 | 385 | 72928 | 2.9 |
| **ERV_classII** | 79 | 16387 | 0.44 | 39 | 8458 | 0.34 |
| **DNA elements:** | 696 | 96124 | 2.6 | 501 | 68906 | 2.74 |
| **hAT-Charlie** | 329 | 41496 | 1.12 | 230 | 30269 | 1.2 |
| **TcMar-Tigger** | 216 | 34349 | 0.93 | 166 | 25275 | 1.01 |
| **Unclassified:** | 28 | 2292 | 0.06 | 19 | 1721 | 0.07 |
| **Total interspersed repeats:** | | 1732973 | 46.93 | | 1151877 | 45.82 |
| **Small RNA:** | 25 | 1893 | 0.05 | 15 | 1247 | 0.05 |
| **Satellites:** | 842 | 187698 | 5.08 | 474 | 103459 | 4.12 |
| **Simple repeats:** | 878 | 35996 | 0.97 | 585 | 23688 | 0.94 |
| **Low complexity:** | 107 | 4911 | 0.13 | 69 | 3201 | 0.13 |
