## Supplementary material for "Chimeric chromosome landscapes of human somatic cell cultures show dependence on stress and regulation of genomic repeats by CGGBP1": Table S10

| **Samples** | **GM02639-CT** | **GM02639-KD** |
|---|---|---|
| Total chimeric reads | 44026 | 19734 |
| Reads with X-U-A chimeras | 2206 | 941 |
| Reads with Y-U-A chimeras | 3046 | 1848 |
| X-U-A (%) | 5.01 | 4.77 |
| Y-U-A (%) | 6.92 | 9.36 |
