## Supplementary material for "Chimeric chromosome landscapes of human somatic cell cultures show dependence on stress and regulation of genomic repeats by CGGBP1": Table S11

| Sample Name | GM01391-CT | GM01391-KD |
| --- | --- | --- |
| Read count | 104768797 | 117629570 |
| Base count | 15447113744 | 17535893169 |
| Mean read length | 147.44 | 149.08 |
| Reads mapped by bowtie2 | 67725152 | 80980111 |
| % mapped reads | 64.64 | 68.84 |
