## Supplementary material for "Chimeric chromosome landscapes of human somatic cell cultures show dependence on stress and regulation of genomic repeats by CGGBP1": Table S12

| **Samples** | **GM01391-CT** | **GM01391-KD** |
|---|---|---|
| Read counts with unexpected allelic identities | 6799763 | 2354827 |
| Read counts with expected allelic identities | 11509946 | 4471689 |
| Somatic mutation rate | 59.08 | 52.66 |
