## Supplementary material for "Chimeric chromosome landscapes of human somatic cell cultures show dependence on stress and regulation of genomic repeats by CGGBP1": Table S13

| Samples | GM02639-CT | GM02639-KD | GM01391-CT | GM01391-KD |
|---|---|---|---|---|
| Reads with maternal allelic identity | 816032 | 394787 | 2053554 | 940277 |
| Reads with paternal allelic identity | 985983 | 466003 | 2376579 | 1002715 |
| Interallelic chimeric read count | 76874 | 39370 | 75879 | 108513 |
| Reads subjected to allelic identification | 1878889 | 900160 | 4506012 | 2051505 |
| Non-chimeric reads per interallelic chimeric event | 23.4 | 21.9 | 58.4 | 17.9 |
