## Supplementary material for "Chimeric chromosome landscapes of human somatic cell cultures show dependence on stress and regulation of genomic repeats by CGGBP1": Table S14

| **Samples** | **Reads with interallelic chimera (without normalization)** | **Reads with interallelic chimera (after normalization)** |
|---|---:|---:|
| **GM02639-CT** | 76874 | 37561 |
| **GM02639-KD** | 39370 | 39370 |
| **GM01391-CT** | 75879 | 75879 |
| **GM01391-KD** | 108513 | 90751 |
