## Supplementary material for "Chimeric chromosome landscapes of human somatic cell cultures show dependence on stress and regulation of genomic repeats by CGGBP1": Table S15

| Repeated measures ANOVA summary | Repeat-free (Fig 2D) | | | | | Alu-SINEs (Fig 2E) | | | | | LINEs (Fig 2F) | | | | | Simple repeats (Fig 2G) | | | | |
|---|---|---|---|---|---|---|---|---|---|---|---|---|---|---|---|---|---|---|---|---|
| Assume sphericity? | Yes | | | | | Yes | | | | | Yes | | | | | Yes | | | | |
| F | 88.9 | | | | | 21.85 | | | | | 22.95 | | | | | 17.59 | | | | |
| P value | <0.0001 | | | | | <0.0001 | | | | | <0.0001 | | | | | <0.0001 | | | | |
| P value summary | **** | | | | | **** | | | | | **** | | | | | **** | | | | |
| Statistically significant (P < 0.05)? | Yes | | | | | Yes | | | | | Yes | | | | | Yes | | | | |
| R squared | 0.6897 | | | | | 0.3533 | | | | | 0.3646 | | | | | 0.3054 | | | | |
| Was the matching effective? | | | | | | | | | | | | | | | | | | | | |
| F | 103.6 | | | | | 32.55 | | | | | 40.59 | | | | | 17.95 | | | | |
| P value | <0.0001 | | | | | <0.0001 | | | | | <0.0001 | | | | | <0.0001 | | | | |
| P value summary | **** | | | | | **** | | | | | **** | | | | | **** | | | | |
| Is there significant matching (P < 0.05)? | Yes | | | | | Yes | | | | | Yes | | | | | Yes | | | | |
| R squared | 0.8654 | | | | | 0.808 | | | | | 0.8376 | | | | | 0.7137 | | | | |
| ANOVA table | SS | DF | MS | F (DFn, DFd) | P value | SS | DF | MS | F (DFn, DFd) | P value | SS | DF | MS | F (DFn, DFd) | P value | SS | DF | MS | F (DFn, DFd) | P value |
| Treatment (between columns) | 4.379 | 5 | 0.8759 | F (5, 200) = 88.90 | P<0.0001 | 3.23 | 5 | 0.6461 | F (5, 200) = 21.85 | P<0.0001 | 2.253 | 5 | 0.4505 | F (5, 200) = 22.95 | P<0.0001 | 4.614 | 5 | 0.9228 | F (5, 200) = 17.59 | P<0.0001 |
| Individual (between rows) | 40.81 | 40 | 1.02 | F (40, 200) = 103.6 | P<0.0001 | 38.49 | 40 | 0.9623 | F (40, 200) = 32.55 | P<0.0001 | 31.86 | 40 | 0.7965 | F (40, 200) = 40.59 | P<0.0001 | 37.67 | 40 | 0.9417 | F (40, 200) = 17.95 | P<0.0001 |
| Residual (random) | 1.97 | 200 | 0.009852 | | | 5.913 | 200 | 0.02957 | | | 3.925 | 200 | 0.01963 | | | 10.49 | 200 | 0.05247 | | |
| Total | 47.16 | 245 | | | | 47.64 | 245 | | | | 38.04 | 245 | | | | 52.78 | 245 | | | |
| Data summary | | | | | | | | | | | | | | | | | | | | |
| Number of treatments (columns) | 6 | | | | | 6 | | | | | 6 | | | | | 6 | | | | |
| Number of subjects (rows) | 41 | | | | | 41 | | | | | 41 | | | | | 41 | | | | |
| Number of missing values | 0 | | | | | 0 | | | | | 0 | | | | | 0 | | | | |
