## Supplementary material for "Chimeric chromosome landscapes of human somatic cell cultures show dependence on stress and regulation of genomic repeats by CGGBP1": Table S16

| Sample Name | HEK293T-CT | HEK293T-KD |
|---|---|---|
| Read count | 90638165 | 83117053 |
| Base count | 13866969515 | 12314145819 |
| Mean read length | 152.99 | 148.15 |
| Reads mapped by bowtie2 | 49198728 | 31046804 |
| Reads remained unmapped | 41439437 | 52070249 |
| Base count for unmapped reads | 5938568698 | 7395489494 |
| Total A-U-B chimeric events in unmapped reads | 4888 | 5472 |
| Chimeric DNA events per billion bases sequenced | 823.09 | 739.91 |
