## Supplementary material for "Chimeric chromosome landscapes of human somatic cell cultures show dependence on stress and regulation of genomic repeats by CGGBP1": Table S17

| Repeat types | HEK293T-CT | | | HEK293T-KD | | |
|---|---|---|---|---|---|---|
| | Number of elements | Length occupied (bp) | Sequence (%) | Number of elements | Length occupied (bp) | Sequence (%) |
| **SINEs:** | 2563 | 381682 | 43.21 | 1992 | 297479 | 29.63 |
| **ALUs** | 2546 | 380050 | 43.02 | 1938 | 291328 | 29.02 |
| **MIRs** | 17 | 1632 | 0.18 | 29 | 4263 | 0.42 |
| **LINEs:** | 740 | 92899 | 10.52 | 1046 | 126105 | 12.56 |
| **LINE1** | 732 | 92378 | 10.46 | 987 | 120840 | 12.04 |
| **LINE2** | 8 | 521 | 0.06 | 57 | 5087 | 0.51 |
| **L3/CR1** | 0 | 0 | 0 | 0 | 0 | 0 |
| **LTR elements:** | 215 | 35041 | 3.97 | 525 | 57657 | 5.74 |
| **ERVL** | 3 | 486 | 0.06 | 76 | 7344 | 0.73 |
| **ERVL-MaLRs** | 161 | 26678 | 3.02 | 221 | 24152 | 2.41 |
| **ERV_classI** | 42 | 6454 | 0.73 | 219 | 24633 | 2.45 |
| **ERV_classII** | 9 | 1423 | 0.16 | 7 | 1194 | 0.12 |
| **DNA elements:** | 51 | 3845 | 0.44 | 83 | 6308 | 0.63 |
| **hAT-Charlie** | 5 | 540 | 0.06 | 29 | 2203 | 0.22 |
| **TcMar-Tigger** | 23 | 1761 | 0.2 | 38 | 3100 | 0.31 |
| **Unclassified:** | 40 | 3089 | 0.35 | 82 | 4828 | 0.48 |
| **Total interspersed repeats:** | | 516556 | 58.48 | | 492377 | 49.05 |
| **Small RNA:** | 149 | 8425 | 0.95 | 112 | 7407 | 0.74 |
| **Satellites:** | 987 | 171056 | 19.36 | 1035 | 160049 | 15.94 |
| **Simple repeats:** | 757 | 62855 | 7.12 | 520 | 44163 | 4.4 |
| **Low complexity:** | 12 | 718 | 0.08 | 7 | 353 | 0.04 |
